# *Wolbachia* strengthens the match between pre-mating and early post-mating isolation in spider mites

**DOI:** 10.1101/2024.05.09.593295

**Authors:** Miguel A. Cruz, Sara Magalhães, Murat Bakırdöven, Flore Zélé

## Abstract

Endosymbiotic reproductive manipulators are widely studied as sources of post-zygotic isolation in arthropods, but their effect on pre-zygotic isolation between genetically differentiated populations has garnered less attention. We tested this using two partially isolated populations of the red and green colour forms of *Tetranychus urticae*, either uninfected or infected with different *Wolbachia* strains, one inducing cytoplasmic incompatibility and the other not. We first investigated male and female preferences, and found that, in absence of infection, females were not choosy but all males preferred red-form females. *Wolbachia* effects were more subtle, with only the CI-inducing strain slightly strengthening colour-form based preferences. We then performed a double-mating experiment to test how incompatible matings affect subsequent mating behaviour and offspring production, as compared to compatible matings. Females mated with an incompatible male (infected and/or heterotypic) were more attractive and/or receptive to subsequent (compatible) matings, although analyses of offspring production revealed no clear benefit for this re-mating behaviour (*i.e.*, apparently unaltered first male sperm precedence). Finally, by computing the relative contributions of each reproductive barrier to total isolation, we showed that pre-mating isolation matches both host-associated and *Wolbachia*-induced post-mating isolation, suggesting that *Wolbachia* could contribute to reproductive isolation in this system.

## Introduction

Understanding the evolution of reproductive barriers between taxa has long been a major focus of evolutionary biology (Coyne and Orr 2004). While speciation research has traditionally viewed species divergence as a process inevitably leading to full reproductive isolation (biological species concept; Mayr 1942), recent evidence has shown that partial isolation occurring along the speciation continuum (Stankowski and Ravinet 2021) can be reversible (Taylor et al. 2006; Bhat et al. 2014; Kearns et al. 2018), or may even be selected for in some circumstances (Servedio and Hermisson 2020). Studying population pairs for which reproductive barriers are incomplete is of great value to understand these processes, as it can provide insight into which type of reproductive barrier is more likely to evolve first, then drive the evolution of others (Baack et al. 2015; Lackey and Boughman 2017). On the one hand, late post-zygotic barriers leading to costly hybridization can evolve first (*e.g.*, in allopatry), then promote the evolution of pre- and/or early post-zygotic barriers at secondary contact (*i.e.*, reinforcement following the definition of Coughlan and Matute; 2020; but see Bank et al. 2012). On the other hand, by limiting gene flow, pre-zygotic barriers should lead to faster accumulation of genetic differences between populations in sympatry, thereby promoting the evolution of post-zygotic barriers (*e.g.*, Lackey and Boughman 2017). In addition, previous work suggested that reproductive isolation may be driven not only by the genetics of the organisms themselves but also by their endosymbionts, especially those that directly manipulate the reproduction of their hosts (Duron et al. 2008; Engelstädter and Hurst 2009; Brucker and Bordenstein 2012).

*Wolbachia* is a widespread endosymbiotic bacterium (Weinert et al. 2015) that manipulates its host reproduction in different ways to increase its own transmission (Werren et al. 2008; Engelstädter and Hurst 2009). The most common of such manipulations is cytoplasmic incompatibility (CI), a conditional sterility phenotype that results in embryonic mortality of offspring from crosses between infected males and uninfected females (or females infected with an incompatible strain; Breeuwer and Werren 1990; Shropshire et al. 2020). Although the contribution of *Wolbachia* to post-zygotic isolation has been extensively studied in different systems, its contribution to pre-zygotic isolation (both pre- and post-mating) between hosts has received comparatively less attention (see Shropshire and Bordenstein 2016; Bi and Wang 2020; Kaur et al. 2021), especially when acting alongside host genetic incompatibilities.

Theory predicts that *Wolbachia* could drive reinforcement between undifferentiated host populations (*i.e.*, females may evolve avoidance of incompatible males to escape CI; Champion de Crespigny et al. 2005; Telschow et al. 2005), but empirical studies have produced contrasting results, most of them showing no (or weak) evidence for CI-driven assortative mating (reviewed by Shropshire and Bordenstein 2016; Bi and Wang 2020). Such discrepancy could be explained by uneven abilities of hosts to detect *Wolbachia* infection in their mates (*e.g.*, *Wolbachia* may alter the chemical profiles of some host species only; Richard 2017; Fortin et al. 2018; Schneider et al. 2019), or because avoidance of CI might be more likely when the infection is associated with pre-existing host traits that can be used for mate recognition (Engelstädter and Telschow 2009). If so, CI avoidance should be more commonly found between already differentiated populations. In line with this prediction, the rare studies focusing on genetically differentiated hosts showed that pre-mating isolation was strengthened (possibly even caused) by *Wolbachia* infection (*e.g.*, Jaenike et al. 2006; Koukou et al. 2006; Miller et al. 2010; but see Shoemaker et al. 1999). Finally, *Wolbachia* infection may also drive post-mating pre-zygotic isolation. For instance, *Wolbachia* infection can have deleterious effects on sperm production or transfer (Snook et al. 2000; Lewis et al. 2011), fertilization success (Bruzzese et al. 2021), or effectiveness of re-mating (De Crespigny and Wedell 2006; Champion De Crespigny et al. 2008; Liu et al. 2014; He et al. 2018). However, to our knowledge, no study has specifically disentangled the relative role of *Wolbachia* from that of host genetic factors on different types of post-mating pre-zygotic barriers.

*Tetranychus* spider mites are an excellent system to address the interplay between host-associated and symbiont-induced incompatibilities (Cruz et al. 2021). *Wolbachia* is ubiquitous in this genus (Breeuwer and Jacobs 1996; Gotoh et al. 2003; Xie et al. 2006; Zhang et al. 2013, 2016; Zélé et al. 2018a), and its effects have been widely studied, especially in the two-spotted spider mite *T. urticae*. Natural populations of this species can be infected with highly variable prevalence (ranging from 0 to 100%; Gotoh et al. 2003, 2007; Zhang et al. 2016; Zélé et al. 2018a,b) of different *Wolbachia* strains, mostly belonging to the *Ori* subgroup of supergroup B (Gotoh et al. 2003, 2007; Zhang et al. 2013; Suh et al. 2015; Pina et al. 2020; although some strains belonging to the *Con* subgroup have also been found; Xie et al. 2006). In this host, the bacterium also induces highly variable degrees of different types of CI (from 0 to 100% of either FM- or MD-type CI, which correspond, respectively, to mortality or development as male of fertilized eggs in incompatible crosses; *e.g.*, Breeuwer 1997; Perrot-Minnot et al. 2002; Vala et al. 2002; Gotoh et al. 2003; Suh et al. 2015; Zélé et al. 2020; Wybouw et al. 2022), and has variable effects on pre-mating isolation (Zhao et al. 2013b; Rodrigues et al. 2022; Vala et al. 2004). However, in spider mites, as in many other arthropod species, the contribution of *Wolbachia* to post-mating pre-zygotic isolation has seldom been studied (though see Cooper et al. 2017 for the *Drosophila yakuba* complex), which is at odds with the critical role that this symbiont may play in the speciation processes currently ongoing in this group.

Given the wide and overlapping distribution of many spider mite species (Migeon and Dorkeld 2023), as well as the high variability in genetic distances both between and within species (*e.g.*, Matsuda et al. 2018; Villacis-Perez et al. 2021), spider mites commonly suffer various degrees of reproductive isolation. In particular, there is ample evidence of variation in all possible post-zygotic reproductive barriers (zygote and juvenile hybrid mortality, hybrid sterility, hybrid breakdown), both between (Keh 1952; Helle and Van de Bund 1962; Hill and O’Donnell 1991) and within spider mite species (*e.g.*, Van de Bund and Helle 1960; de Boer 1982a,b; Sugasawa et al. 2002; Knegt et al. 2017; Cruz et al. 2021). Several studies also revealed variable post-mating pre-zygotic isolation in this group (*e.g.*, fertilization failure resulting from gametic or mechanical incompatibilities), as evidenced by a reduction in the production of female offspring, given that spider mites are arrhenotokous haplodiploids (haploid males develop from unfertilized eggs and diploid females from fertilized eggs; Helle and Bolland 1967). Hence, whereas no female offspring are produced in crosses between well-formed species (*e.g.*, Helle and Van de Bund 1962; Hill and O’Donnell 1991; Chain-Ing and Sheuan-Ping 1995; Clemente et al. 2016, 2018), male-biased sex ratios are often reported in crosses between genetically differentiated ‘forms’ of the same species or even between genotypes of the same form (*e.g.*, Gotoh et al. 1993; Navajas et al. 2000; Sugasawa et al. 2002; Auger et al. 2013; Cruz et al. 2021; Villacis-Perez et al. 2021). In addition, because spider mites exhibit first male sperm precedence (only the first male that mates with a female sires all the offspring; Helle 1967; Rodrigues et al. 2020), females usually cannot restore their fitness through re-mating. Therefore, post-mating incompatibilities are particularly costly and should select for earlier pre-zygotic barriers through reinforcement. Yet, highly variable degrees of pre-mating isolation can be found both between (Sato et al. 2014, 2016; Clemente et al. 2016; Sato and Alba 2020) and within species (*e.g.*, Murtaugh and Wrensch 1978; Gotoh et al. 1993).

To improve our understanding of the contributions that *Wolbachia* can make to reproductive isolation among its hosts, we aimed at disentangling the relative contributions of *Wolbachia* and host genetic factors to the strength of both pre- and post-mating pre-zygotic barriers between two closely-related colour forms, green and red, of the two spotted spider mite *T. urticae*, which are sometimes also referred to as separate species: *T. urticae* and *T. cinnabarinus* (Xie et al. 2006; Auger et al. 2013; Lu et al. 2017, 2018). Indeed, although a recent study showed very high differentiation between populations of these two forms at the genomic level (Xue et al. 2023), post-zygotic isolation was found to range from full to only partial (Murtaugh and Wrensch 1978; Dupont 1979; de Boer 1982a,b; Sugasawa et al. 2002), and they do not seem to differ in the prevalence or type of *Wolbachia* strains they carry, nor, overall, in CI level or pattern they induce (*e.g.*, Perrot-Minnot et al. 2002; Vala et al. 2002; Xie et al. 2006; Gotoh et al. 2007; Zélé et al. 2020; Wybouw et al. 2022). Moreover, the two forms have an overlapping worldwide distribution (Hinomoto et al. 2001; Lu et al. 2017; Godinho et al. 2020; Migeon and Dorkeld 2023; Xue et al. 2023), they share the same host plant range (Auger et al. 2013) and sometimes the same individual plant (Lu et al. 2017; Zélé et al. 2018b). A previous study focusing on the joint effects of *Wolbachia*-induced and host-associated post-mating incompatibilities between populations of these two forms revealed full reproductive isolation due to late post-zygotic barriers (hybrid sterility and hybrid breakdown) that were independent of *Wolbachia* infection (Cruz et al. 2021). However, this study also revealed partial and asymmetrical earlier post-mating barriers (pre- and/or post-zygotic), resulting from a combination of host-associated and *Wolbachia*-induced incompatibilities (Cruz et al. 2021). Host genetic incompatibilities led to an increased proportion of haploid sons in detriment of diploid daughters (‘male development’ or MD-type incompatibility, likely due to fertilization failure) in crosses between red-form males and green-form females, whereas the reciprocal cross yields no change in sex ratio. Moreover, whereas *Wolbachia* infection in green-form males was not associated with CI induction of any type, *Wolbachia* infection in red-form males led to an increased embryonic mortality of their daughters (‘female mortality’ or FM-type CI). Furthermore, both types of incompatibility had additive effects and acted in the same direction of crosses (Cruz et al. 2021), which hinted at a possible role of *Wolbachia*-induced incompatibilities in host population divergence and subsequent evolution of intrinsic reproductive barriers, as found in *Nasonia* wasps (Bordenstein et al. 2001).

Here, we significantly build upon previous work by investigating pre- and post-mating pre-zygotic reproductive barriers between these spider mite populations. We first performed male and female choice tests to determine preference for infected or uninfected mates from different colour-form populations (*i.e.*, test for pre-mating isolation; Table 1). Second, we used a no-choice test to investigate the effect of female mating history (virgin or previously-mated with a compatible *vs* incompatible male) on mating behaviour, and to test whether eggs are more likely fertilized by compatible than incompatible sperm (*i.e.*, test for ‘homotypic’ sperm precedence; Table 2). Finally, we used data gathered throughout all experiments stemming from this study and the previous one (Cruz et al. 2021) to estimate the relative contribution of each measured host-associated or *Wolbachia*-induced individual barrier to total reproductive isolation in this system.

**Table 1.**
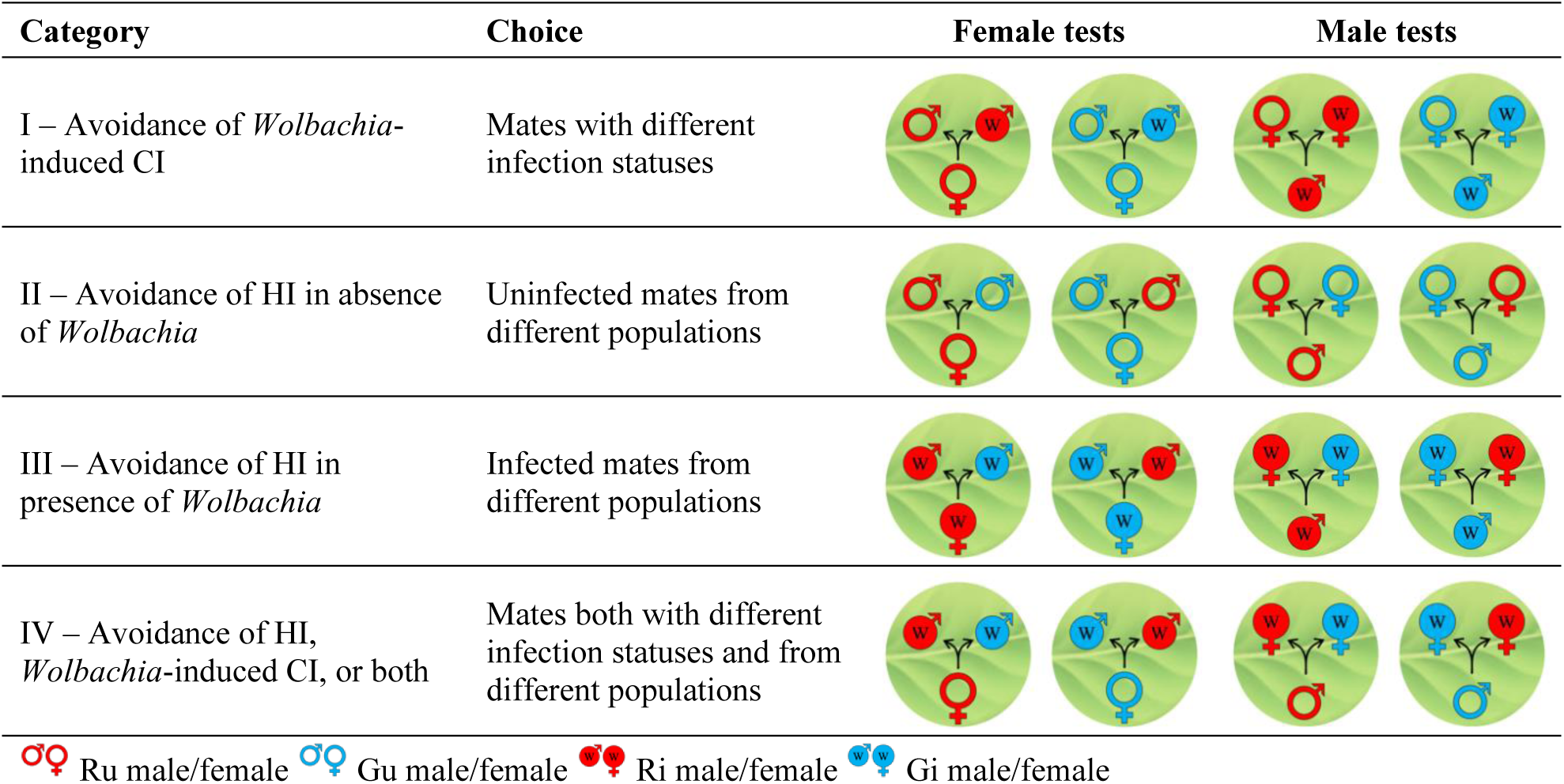
Choice tests to assess the mating behaviour and preference of males or females that were given the choice between two mates of different colour forms and/or *Wolbachia* infection status. CI: cytoplasmic incompatibility; HI: host-associated incompatibility.

**Table 2.**
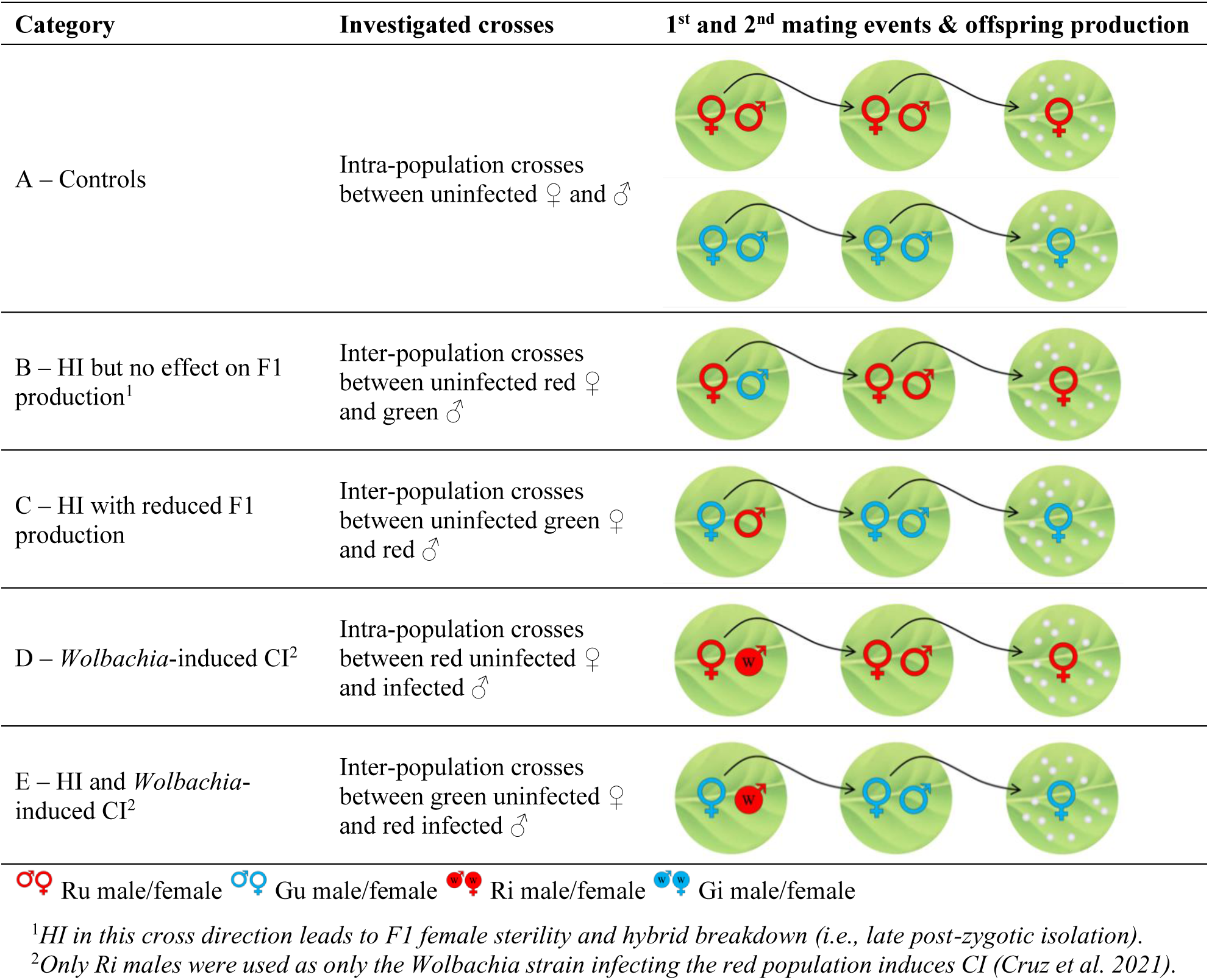
No-choice tests to assess the behaviour and offspring production of virgin females (♀) placed with a compatible or incompatible male (of a similar or different colour form and/or *Wolbachia* infection status, respectively; 1^st^ mating event), then with a second compatible male from their own population (only for females that had mated with the first male; 2^nd^ mating event). CI: cytoplasmic incompatibility; HI: host-associated incompatibility.

## Materials and Methods

### Spider mite populations

Two populations of spider mites, each belonging to a different colour form of *T. urticae* (‘red’ or ‘green’), and either infected or uninfected with *Wolbachia*, were used in this study. These populations, fully described in the Supplementary Box S1, were previously used to assess post-mating isolation caused by both host-associated incompatibilities (HI) and *Wolbachia*-induced reproductive barriers (Cruz et al. 2021). Briefly, the *Wolbachia*-infected population ‘Ri’ and its uninfected counterpart ‘Ru’ (‘Ri1’ and ‘Ru1’ in Cruz et al. 2021) belong to the red form of *T. urticae*, whereas the *Wolbachia*-infected population ‘Gi’ and the uninfected population that derived from it, ‘Gu’, belong to the green form of *T. urticae*. The original Ri and Gi populations, both fully and stably infected with *Wolbachia* at the time of the experiments, were collected from locations *ca.* 34 km apart in the region of Lisbon (Portugal; see Box S1). In this region, the prevalence of *Wolbachia* is very high in red form populations (40 to 100%, *ca*. 94% on average; Zélé et al. 2018a,b) and CI levels are moderate to high (*ca*. 27 to 65%; Zélé et al. 2020), as for the Ri population used here (naturally fully infected and *ca*. 57 ± 3% CI; Zélé et al. 2018a, 2020). The incidence of *Wolbachia* in green form populations of this same region seems comparatively much lower (only 1 out of 4 populations were found carrying the symbiont, as reported in Godinho et al. 2020), perhaps due to weak or no CI induction as previously found for the Gu population used here (unknown prevalence but no CI induction; Cruz et al. 2021). The Ru and Gu populations were then obtained from antibiotic treatments as detailed in Box S1. All populations were reared at high numbers (>1000 females per population) in mite-proof cages containing bean plants (*Phaseolus vulgaris*, cv. Contender seedlings obtained from Germisem, Oliveira do Hospital, Portugal) under the same standard laboratory conditions (24±2°C, 16/8h L/D). All behavioural observations were conducted during daytime at constant room temperature (25 ± 2°C).

### Mate preference and behaviour of males and females in choice tests

To determine whether spider mites discriminate between mates to avoid *Wolbachia*-induced and/or host-associated incompatibilities, individual males and females were provided two mates from different populations and/or infection statuses. All combinations of choice tests performed are described in Table 1. To obtain a large number of individuals of similar age, age cohorts were created for each population (each cohort was used for two to three sequential days of observation). To this aim, 50 mated females or 50 virgin females (to obtain cohorts of females or males, respectively) from each population laid eggs during 3 days on detached bean leaves placed on water-soaked cotton in petri dishes under standard laboratory conditions (24±2°C, 16/8h L/D). Ten to twelve days later, female and male deutonymphs undergoing their last moulting stage (*i.e.*, teleiochrysalids) were randomly collected from the cohorts and placed separately on bean leaf fragments (*ca.* 9cm^2^) to obtain virgin adult females and males of similar age two days later. As opposed to females, males cannot easily be identified based on their body colouration, hence they were painted before each observation with either blue or white water-based paint (randomized across treatments) using a fine brush. Previous experiments showed no effect of this paint on spider mite mate choice and behaviour (Rodrigues et al. 2017, 2022). Subsequently, a pair of virgin mates was placed on a 0.4cm^2^ leaf disc (called ‘arena’ hereafter), then the observation started when the focal individual was also introduced to the arena. The colour of the mate that first copulated with each focal individual was registered, and later assigned to the corresponding treatment (thus ensuring that the observer was blind to the treatment to which mites belonged). The time until the beginning of copulation (‘latency to copulation’) and its duration (‘copulation duration’) were recorded using an online chronometer (http://online-stopwatch.chronme.com/). Each observation lasted until the end of a first copulation or for 30 minutes if no mating occurred. Male and female choice tests were performed separately, each with one replicate of each treatment observed simultaneously per session and four sessions of observations carried out per day. In total, 60 replicates per treatment were obtained over the course of 15 days for each of the two tests.

### (Re)mating behaviour and offspring production in the no-choice test

#### Mating behaviour in the first mating event

Spider mites may possess pre-zygotic mechanisms other than mate discrimination to avoid and/or reduce the cost of incompatibilities, such as rejecting a mate after a copulation has started. Moreover, copulations lasting less than 30 seconds can be insufficient to fully fertilize a spider mite female (Potter and Wrensch 1978; Satoh et al. 2001), which might explain the excessive production of males to the detriment of females (*i.e.*, arising from unfertilized and fertilized eggs, respectively) previously observed in the brood of green females mated with red males (Cruz et al. 2021). To test whether such post-copulatory mechanisms of avoidance of incompatibilities occur in spider mites, we performed a no-choice test, where the mating behaviour of virgin females placed with a single male was observed. Given the workload involved in this experiment, we only performed the crosses allowing to test for the single and combined effects of host-associated and *Wolbachia*-induced incompatibility, along with their respective controls (see Table 2). Also, because only few individuals could be tested per day, male and female teleiochrysalids were directly sampled from the base populations two days prior to observation, and isolated on bean leaf fragments to ensure their virginity. For each treatment, one male and one female were placed together on a 0.5cm^2^ bean leaf disc and observed for 60 minutes. During that time, multiple mating could occur. Thus, in addition to the mating propensity (*i.e.*, the probability that mating occurred at least once) and latency to the first copulation, the copulation frequency (*i.e.*, the number of copulations during the observation period) and the duration of each copulation, to compute the cumulative copulation duration of each couple, were also recorded. At the end of the observation period, females for which at least one copulation occurred were individually placed on a 2cm^2^ bean leaf disc and kept for the next step (see below), while non-mated females and all males were discarded.

#### Mating behaviour in the second mating event

In species with first male sperm precedence such as *T. urticae*, females usually have low receptivity to a second mate (Clemente et al. 2016). However, if the first copulation is interrupted or (at least partially) ineffective, females may show increased receptivity to second matings that could effectively contribute to fertilization (Helle 1967; Clemente et al. 2016; Costa et al. 2023). To test this, females for which at least one copulation occurred during the first mating event were placed with a second compatible male 24 hours later (see Table 2) and their mating behaviour was recorded for 60 minutes as in the first mating event. At the end of the observation period, males were discarded and females were kept individually on 2 cm^2^ bean leaf discs placed on water-soaked cotton in petri dishes in a climatic chamber (25 ± 2°C, 60% RH, 16/8 h L/D).

#### Offspring production and strength of post-mating incompatibilities

To test whether the second copulation with a compatible male could restore female offspring production, the offspring produced over 3 days of oviposition by females mated with either a single or two different males was compared, and female mortality during that period also registered. The number of unhatched eggs was counted 6 days later (day 9), and the numbers of dead juveniles, adult males and females were counted 3 and 6 days later (days 12 and 15). Then, to determine the proportion of offspring affected by host-associated MD-type incompatibility (*i.e.*, “Male Development”), and/or *Wolbachia*-induced FM-type incompatibility (*i.e.*, “Female Mortality”), we computed two indices as fully described in Cruz et al. (2021): the *MD_corr_* index, which calculates the overproduction of sons in the brood (using the number of adult sons and the total number of offspring), and the *FM_corr_* index, which calculates the embryonic mortality of fertilized offspring (using the number of unhatched eggs and the number of adult daughters), both relative to the control crosses to account for background variation. Hence, higher values of *MD_corr_* indicate a greater proportion of sons in the brood to the detriment of daughters, while higher values of *FM_corr_* indicate a greater mortality of female embryos. Finally, as in Cruz et al. (2021), we also computed the proportion of F1 females over the total number of eggs (FP) to determine the combined effect of FM- and MD-type incompatibilities on the total proportion of daughters in each cross. Raw data are shown in the Supplementary Figure S1.

Given the workload and the multiplicity of tasks involved in the entire experiment (first and second mating events, as well as offspring production), only 9 couples were observed simultaneously for each mating event, corresponding to one or two replicates per treatment. Four sessions of observation were performed per day (hence 6 replicates of each cross category per day), each day corresponding to an experimental ‘block’. In total, 19 blocks, each separated by 3 days, were performed to obtain *ca.* 100 replicates per cross category (regardless of whether the females mated during the first and/or second mating events).

### Strength and contribution of each reproductive barrier to total isolation

#### Strength of reproductive isolation for each reproductive barrier (RI_n_)

To estimate the strength of pre- or post-mating reproductive barriers identified for each type of cross within and between the green- and red-form populations, we used the pre-mating data obtained here and the post-mating data from Cruz et al. (2021), respectively. Only reproductive barriers found to play a role in reducing gene flow among the spider mite populations were considered (see. Figure S2): mate preference (*RI_1_*), fertilization failure (*RI_2_*), hybrid inviability (*RI_3_*), hybrid sterility (*RI_4_*), and hybrid breakdown (*RI_5_*), with homotypic sperm precedence and female choice not being included (see Results).

To determine the strength of pre-mating isolation (*RI_1_*), we applied a sexual isolation index, which varies between zero and one, to the male choice data. This index, adapted from Bateman (1949) and Merrell (1950) by Malogolowkin-Cohen et al. (1965), is given by:

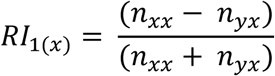

where *n_xx_* is the number of copulations observed between females and males of a population *x*, and *n_yx_* is the number of copulations observed between females of a population *y* and males of the population *x*. As *RI_1_* represents the degree to which a population *x* is isolated from a population *y* due to mating preferences, it was set to 0 in the case of preference for heterotypic mates (*i.e.,* no negative impact on gene flow).

To determine the strength of post-mating barriers, we used the data from Cruz et al. (2021), as late reproductive barriers (*i.e.*, hybrid sterility and hybrid breakdown) were not measured here. Moreover, earlier post-mating barriers (fertilization failure and hybrid inviability) have been estimated in all possible crosses and more precisely in the previous study that focused specifically on post-mating isolation (*i.e.*, experiments were done with larger sample sizes, as they only included single mating treatments). For fertilization failure (*RI_2_*) and hybrid inviability (*RI_3_*) we used the median values of the *MD_corr_* and *FM_corr_* indices in Cruz et al. (2021), which correspond to the percent increase in non-fertilized eggs and in embryonic mortality of fertilized eggs, respectively (see *‘Offspring production and strength of post-mating incompatibilities*’ above). For hybrid sterility (*RI_4_*) and hybrid breakdown (*RI_5_*), we computed the percent decrease in ovipositing F1 females and increase in embryonic mortality of F1 females’ eggs relative to compatible crosses, respectively.

#### Contribution of each reproductive barrier (C_n_) to total isolation (T)

We employed a method previously adapted from Coyne and Orr (1989, 1997) by Ramsey and colleagues (2003), in which total (cumulative) reproductive isolation between two populations or species is computed as a multiplicative function of the strength of each reproductive barrier (*RI_n_*; see above), so that the contribution of each barrier to reducing gene flow at a stage *n* in life history is calculated as:

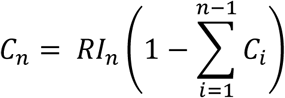

Thus, a given reproductive barrier eliminates gene flow that has not been prevented by earlier barriers, and for *m* reproductive barriers, total reproductive isolation is given by:

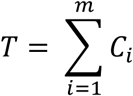

### Statistical analyses

All analyses were carried out using Mixed Models with the R statistical software (v3.6.1). The general procedure used for building all statistical models (R packages, datasets and sample size, coding of the response variables, choice of error structures, usage of random vs. fixed explanatory variables and their interactions); for the simplification of maximal models (containing the complete set of explanatory variables) into minimal models to establish the significance of the explanatory variables (Crawley 2007); and to determine significant changes relative to the intercept as well as to perform contrasts analyses between factor levels, was always substantially the same. It is fully detailed in supplementary materials Box S2, and Tables S1 and S2.

## Results

### Male and female mating behaviour in choice tests

Overall, the propensity of females to mate with one of two provided males depended on their own population (‘focal’: χ^2^_3_=10.06, *p*=0.018; Model 1.1 in Table S1; Figure 1a), with green females being, on average, *ca.* 20% less likely to mate than red females (Tables S3 and S4). However, their mating propensity was unaffected by the type of males they were offered (‘mates’: χ^2^_3_=2.70, *p*=0.44; Model 1.1), and none of them showed any clear mating preference (‘focal’: χ^2^_4_=2.91, *p*=0.57, and ‘mates’: χ^2^_4_=3.47, *p*=0.48; Model 1.2; Figure 1b and Table S5). Conversely, the mating propensity and the mate choice of males were independent of their population (‘focal’: χ^2^_3_=4.13, *p*=0.25 and χ^2^_4_=5.01, *p*=0.29 in Model 1.5 and 1.6, respectively), but strongly affected by the type of females provided (‘mates’: χ^2^_3_=15.72, *p*=0.001 and χ^2^_4_=50.24, *p*<0.0001 in Model 1.5 and 1.6, respectively; Figures 1c and 1d). Indeed, males that were given the choice between two green females were less likely to mate than those that faced a choice that involved a red female (*ca.* 40% *vs* 68% mated males on average; Figure 1c, Tables S3 and S4), and males of either colour form showed a preference for red females (*ca.* 60% to 80% preference; Tables S3 and S4).

**Figure 1.**
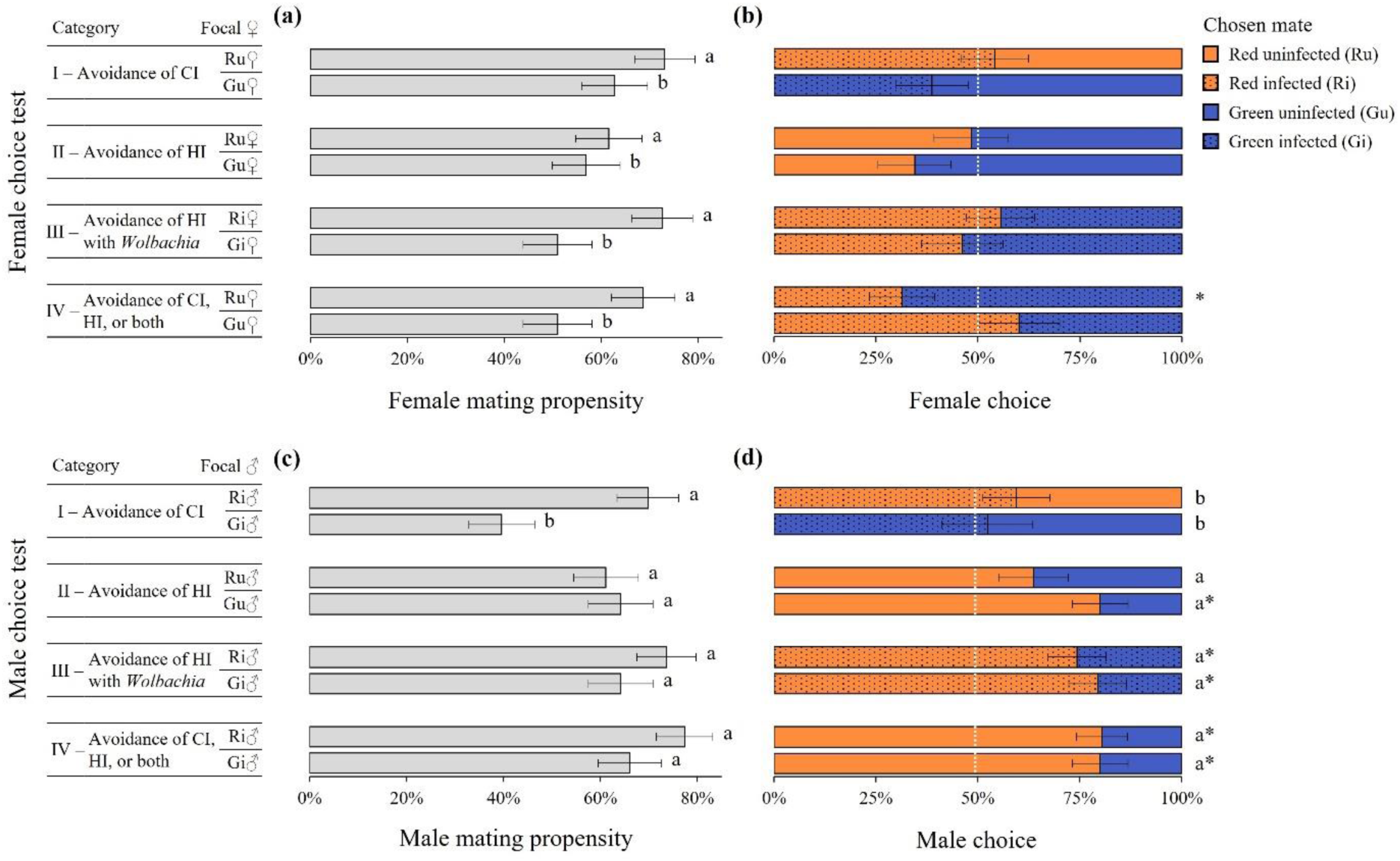
Mating propensity and mate choice of spider mites of different colour forms and/or *Wolbachia* infection status. For each type of choice test, bars represent the mean (± s.e.) proportion of mated (a) females and (c) males, and of mates chosen by (b) females and (d) males (dotted: *Wolbachia*-infected mates; plain: uninfected mates; orange: red mates; blue: green mates). Identical or absent superscripts indicate non-significant differences at the 5% level among treatments (see Table S4), and asterisks indicate a difference to random mating (white dotted line; see Table S5). CI: cytoplasmic incompatibility; HI: host-associated incompatibility.

In contrast to the mite colour form, *Wolbachia* infection had no effect on mite mating propensity and only a small effect on their mate preference (see Figure 1, Table S3 and contrasts in Table S4). Neither uninfected females nor infected males showed any preference between infected or uninfected mates of the same colour (see category I in Figures 1b and 1d), but *Wolbachia* infection in red males (CI-inducing *Wolbachia* strain; Cruz et al. 2021) strengthened their preferences for red females. Indeed, although the mate preference of Ru and Ri males did not differ significantly, Ru males showed no significant difference from random mating (see category II in Figure 1d) whereas Ri males significantly preferred red females over green females (see categories III and IV; see also Table S5). In addition, whereas red females showed no preference between males of either colour form when these were from the same infection status as themselves (see categories II and III in Figure 1b), Ru females preferentially mated with Gi males over Ri males, suggesting avoidance of the CI induced by *Wolbachia* infection in red males (see category IV; see also Table S5). Conversely, the non-CI-inducing *Wolbachia* strain infecting green males (Cruz et al. 2021) had no effect on the strength of mate preference of both males and females.

Finally, latencies to copulation did not differ significantly among focal females or chosen males in the female choice test (χ^2^_3_=6.76, *p*=0.08 and χ^2^_3_=1.35, *p*=0.72, respectively; Model 1.3), nor among focal males or chosen females in the male choice test (χ^2^_3_=1.33, *p*=0.72 and χ^2^_3_=1.03, *p*=0.79, respectively; Model 1.7, Figure 2a,b), but copulation duration differed between males of different colours (Figure 2c) and between females of different infection status (Figure 2d). Regardless of *Wolbachia* infection (although Ru males showed intermediate copulation durations in the female choice test; Figure 2c; Table S6 and S4), green males copulated on average 37 and 40 seconds longer than red males in the female and male choice test, respectively (‘chosen’: χ^2^_3_=7.92, *p*=0.048, and ‘focal’: χ^2^_3_=27.09, *p*<0.0001; Model 1.4 and 1.8, respectively). Conversely, the copulation duration of females was not affected by their colour form (although Gi females showed intermediate copulation durations in the female choice test; Figure 2d and Table S6; see contrasts in Table S4), but that of infected females was, on average, *ca.* 29 and 34 seconds shorter than that of uninfected females in the female and male choice test, respectively (‘focal’: χ^2^_3_=10.64, *p*=0.014, and ‘chosen’: χ^2^_3_=24.70, *p*<0.0001 in Model 1.4 and 1.8, respectively).

**Figure 2.**
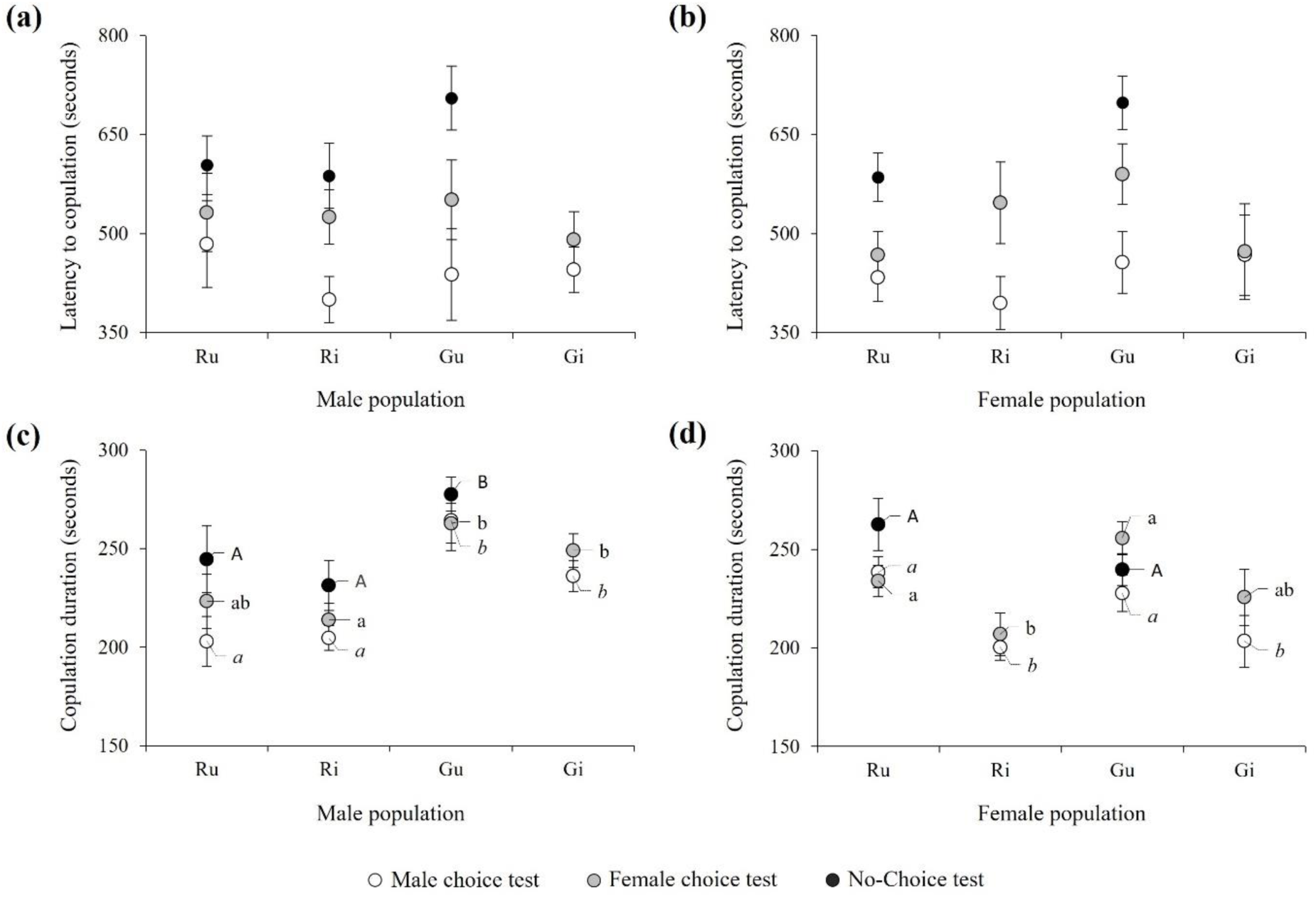
Latency to copulation (a, b) and copulation duration (c, d) of virgin mites during the choice and no-choice tests. Dots represent mean time (± s.e.) in seconds observed for males (a, c) and females (b, d) in the male and female choice tests (white and grey dots, respectively), and in the first mating event of the no-choice test (black dots), regardless of the identity of their mate. The panels (a, c) thus display results obtained for focal males in the male choice test, or male mates in the female choice and no-choice tests, whereas panels (b, d) display results obtained for focal females in the female choice and no-choice tests, or female mates in the male choice test. No significant differences between latencies to copulation were found among the different types of males or females in the choice tests (statistical results are not given for the no-choice test as latencies to copulation exceeding 30 minutes were excluded from the means displayed in this figure to allow comparisons across experiments). For copulation duration, identical superscripts indicate non-significant differences at the 5% level within each test (Italic: male choice test, see Table S4; lowercase: female choice test, see Table S4; uppercase: 1^st^ mating event of the no-choice test, see Table S9). Note that infected females were not used in the no-choice test (hence, black dots are not displayed for Ri and Gi females in panel d).

### (Re)mating behaviour in the no-choice test

On average, 58% of the virgin females placed on a leaf disc with a single male mated within 1 hour, whereas less than 20% of those mated females re-mated when placed with another male 24 hours later. In line with this, a reduced copulation frequency (1.6±0.1 *vs* 2.1±0.1 copulations per couple) and copulation duration (118±13 *vs* 252±8 seconds) and an increased latency to copulation (1582±108 *vs* 986±46 seconds), were observed, on average, between the first and second mating events (Figures 3,4 and Table S7). However, this reduction in the willingness to mate varied across types of crosses for all behavioural traits tested except for copulation frequency (no statistically significant differences found among crosses for either or among the two mating events; see models 2.2 to 2.4 in Table S2).

**Figure 3.**
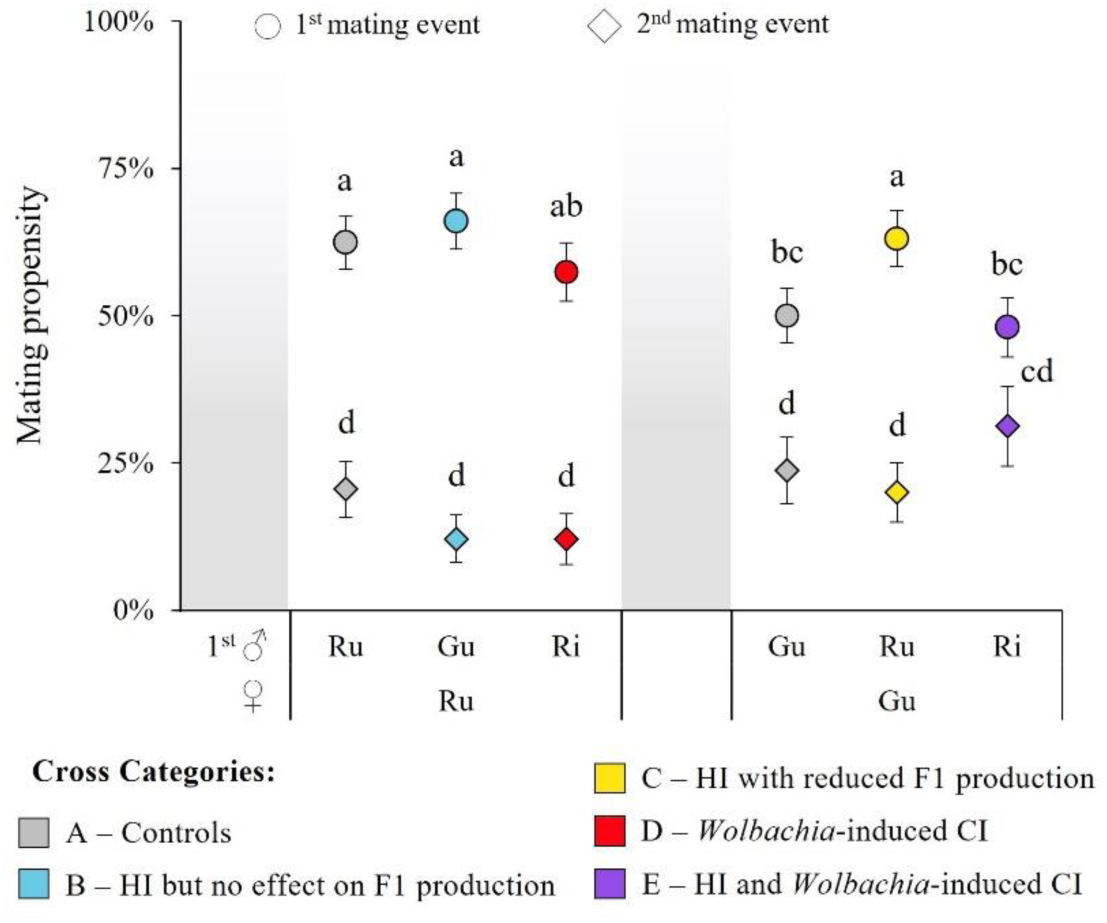
Mating propensity observed in two successive mating events in the no-choice test. For each cross category, circles and diamonds represent mean (± s.e.) proportion of females that mated during the first and the second mating event, respectively. The population of the female is displayed at the bottom level of the x-axis and the population of the first male at the top level (the population of the second male is always the same as that of the female). Identical superscripts indicate non-significant differences at the 5% level among crosses across mating events (see Table S8). CI: cytoplasmic incompatibility; HI: host-associated incompatibility.

**Figure 4.**
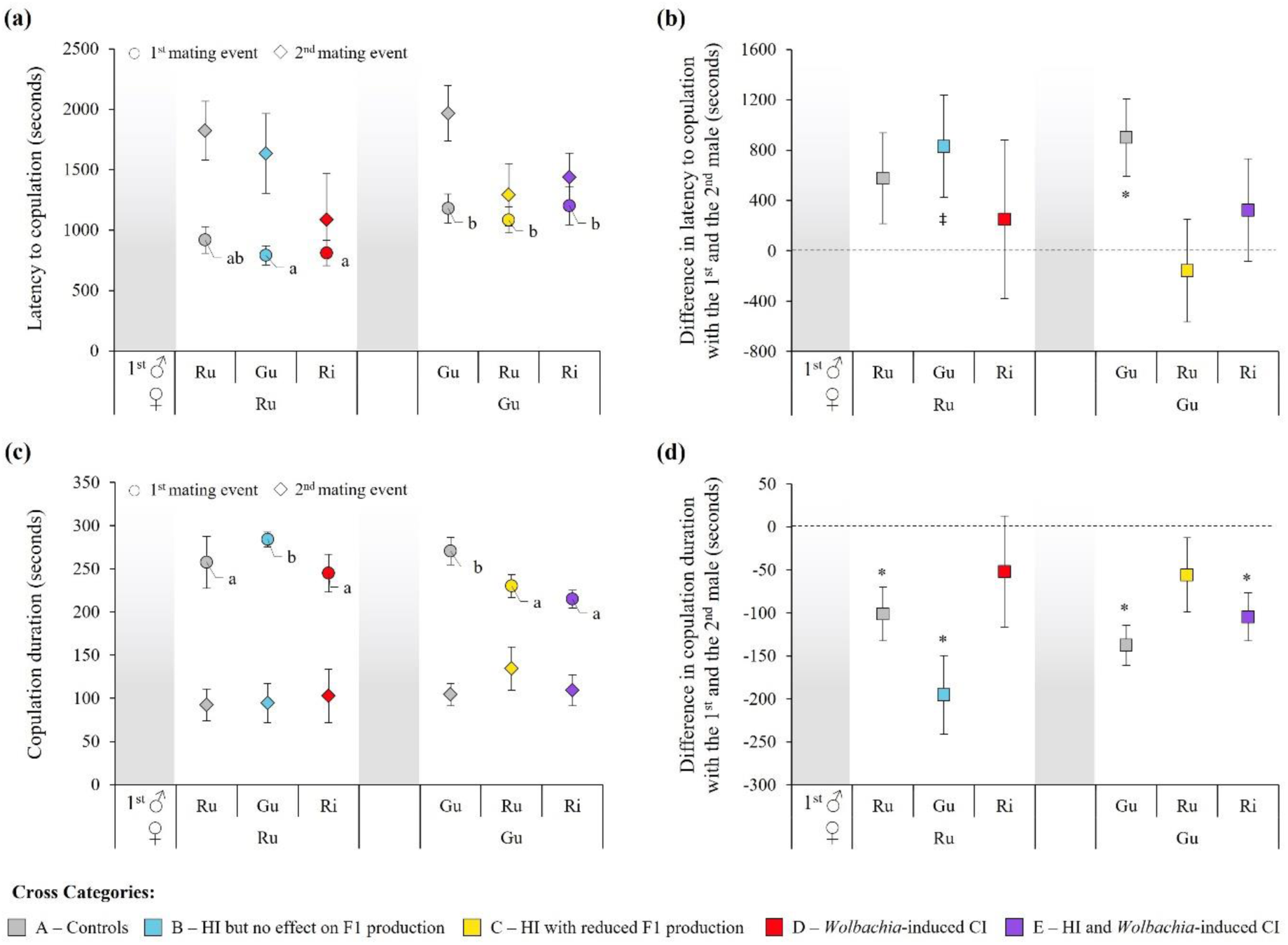
Latency to copulation (a, b) and copulation duration (c, d) observed in two successive mating events in the no-choice test. In (a) and (c), circles and diamonds represent mean time (± s.e.) in seconds for each cross category during the first and the second mating event, respectively. Identical or absent superscripts indicate non-significant differences at the 5% level among crosses within each mating event (see Table S9). In (b) and (d), squares represent the mean time difference (± s.e.) observed between the two mating events for each female that mated with both males (*i.e.* [time spent for the second mating] – [time spent for the first mating]). Superscripts indicate significant differences from zero at the 10% level (‡: *p*<0.10) and at the 5% level (*: *p*≤0.05; see Table S10). In all panels, the population of the female is displayed at the bottom of the x-axis and the population of the first male at the top (the population of the second male is always the same as that of the female). CI: cytoplasmic incompatibility; HI: host-associated incompatibility.

For the mating propensities observed during the first mating event, we found the same tendencies as in the choice tests: Gu females were less likely to mate than Ru females (except when paired with Ru males here), and *Wolbachia* infection in red males seemed to promote mate discrimination (Ri males were less willing to mate with Gu females than Ru males were, whereas both types of males mated as much with Ru females; Figure 3; Tables S7 and S8). Then, although no statistically significant differences were found among crosses in the second event, not all crosses led to the same reduction in mating propensity between the two mating events (*cross x mating event interaction*: χ^2^_5_=16.56, *p*=0.005; model 2.1; Figure 3; Table S8): Gu females showed a lower reduction in their tendency to re-mate than Ru females, especially when they were previously mated with an incompatible Ri male (hence when both types of incompatibilities were at play; Figure 3; Tables S7 and S8).

Conversely to the previous experiment in which virgin individuals could choose their mate and were given only half an hour to mate, we found significant differences among latencies to copulation of couples that mated at least once during the first mating event (χ^2^_5_=13.19, *p*=0.02; model 2.5). Gu females took, on average, 5 more minutes than Ru females to engage in copulation with their first partner, regardless of the form or infection status of the latter (although Ru x Ru crosses had intermediate latencies to copulation; see circles in Figure 4a; Tables S7 and S9). Also, as in the previous experiment (choice tests), the cumulative time spent copulating was longer for green males than for red males regardless of their infection status and the female they mated with (*ca.* 39 seconds difference; χ^2^_5_=21.19, *p*<0.001; model 2.4; Figures 2c,d and 4c; Tables S7 and S9). Then, when females that mated during the first mating event were placed with a second male 24 hours later, their latency to copulation increased by almost 10 minutes, and their copulation duration was more than 2 minutes shorter, than when they were virgin (see diamonds in Figure 4a,c; Table S7). Despite no significant differences being found among types of crosses for both latency to copulation and cumulative copulation duration in the second mating event (χ^2^_5_=5.16, *p*=0.40; model 2.6; Figure 4a, and χ^2^_5_=2.78, *p*=0.73; model 2.9; Figure 4c, respectively), behavioural changes between the two mating events at the female level (for those who mated in both events) varied depending on the type of cross (χ^2^_6_=12.47, *p*=0.05; model 2.7; Figure 4b, and χ^2^_6_=43.73, *p*<0.0001; model 2.10; Figure 4d, for latency to copulation and cumulative copulation duration, respectively). Thus, in line with the mating propensity observations, differences in latency to copulation and copulation duration between mating events tended to disappear for females that had first mated with an incompatible male (except for the copulation duration of Gu females mated with Ri males; Figures 4b,d; Table S10).

### Effect of re-mating on offspring production in the no-choice experiment

The pattern of offspring production for females that mated only with one male (Figure 5a) was consistent with that described in our previous study (Cruz et al. 2021). Briefly, (i) we found an overproduction of males (MD-type incompatibility) in crosses between green females (Gu) and red males (Ru or Ri) as compared to the other crosses (χ^2^_5_=76.30, *p*<0.0001; model 2.12; Figure 5b). Moreover, because copulations were observed in the present study, it further unambiguously revealed a high variability for this barrier: among the 66 Gu females that mated only with a Ru or Ri mate and oviposited (*i.e.*, 85 Gu females mated with a Ru or Ri male subsequently refused to mate with a second male; Table S7, and 19 of these females did not lay a single egg), 20 produced only sons (*i.e.*, full incompatibility), whereas 18 did not produce a more male-biased sex ratio than the controls (*i.e.*, no incompatibility); (ii) we found an increased female embryonic mortality (FM-type CI quantified as a decreased hatching rate of fertilized eggs) in crosses between uninfected females (Gu or Ru) and males infected with a CI-inducing *Wolbachia* strain (Ri males), as compared to the other crosses (χ^2^_5_=76.78, *p*<0.0001; model 2.13; Figure 5c); and (iii) we found a reduction in the proportion of daughters (FP) in crosses affected by either (or both) type(s) of incompatibility (*i.e.*, Ru x Ri, Gu x Ru and Gu x Ri, female x male crosses) as compared to compatible crosses (χ^2^_5_=87.65, *p*<0.0001; model 2.14; Figure 5d). However, no difference in offspring production was found between females that mated with only one or two different males (daily fecundity: χ^2^_1_=3.19, *p*=0.07; model 2.11; MD_corr_: χ^2^_1_=0.63, *p*=0.43; model 2.12; FM_corr_: χ^2^_1_ =0.14, *p*=0.71; model 2.13; FP: χ^2^_1_=0.17, *p*=0.68; model 2.14), regardless of whether the first male was compatible or not (*i.e.*, no significant interactions between the type of cross and whether females mated with one or two males; daily fecundity: χ^2^_5_ =4.57, *p*=0.47; model 2.11; MD : χ^2^_5_ =0.89, *p*=0.97; model 2.12; FM_corr_: χ^2^_5_ =9.33, *p*=0.10; model 2.13; FP: χ^2^_5_=7.17, *p*=0.21; model 2.14; Figure 5; see also Figure S1 and Table S7).

**Figure 5.**
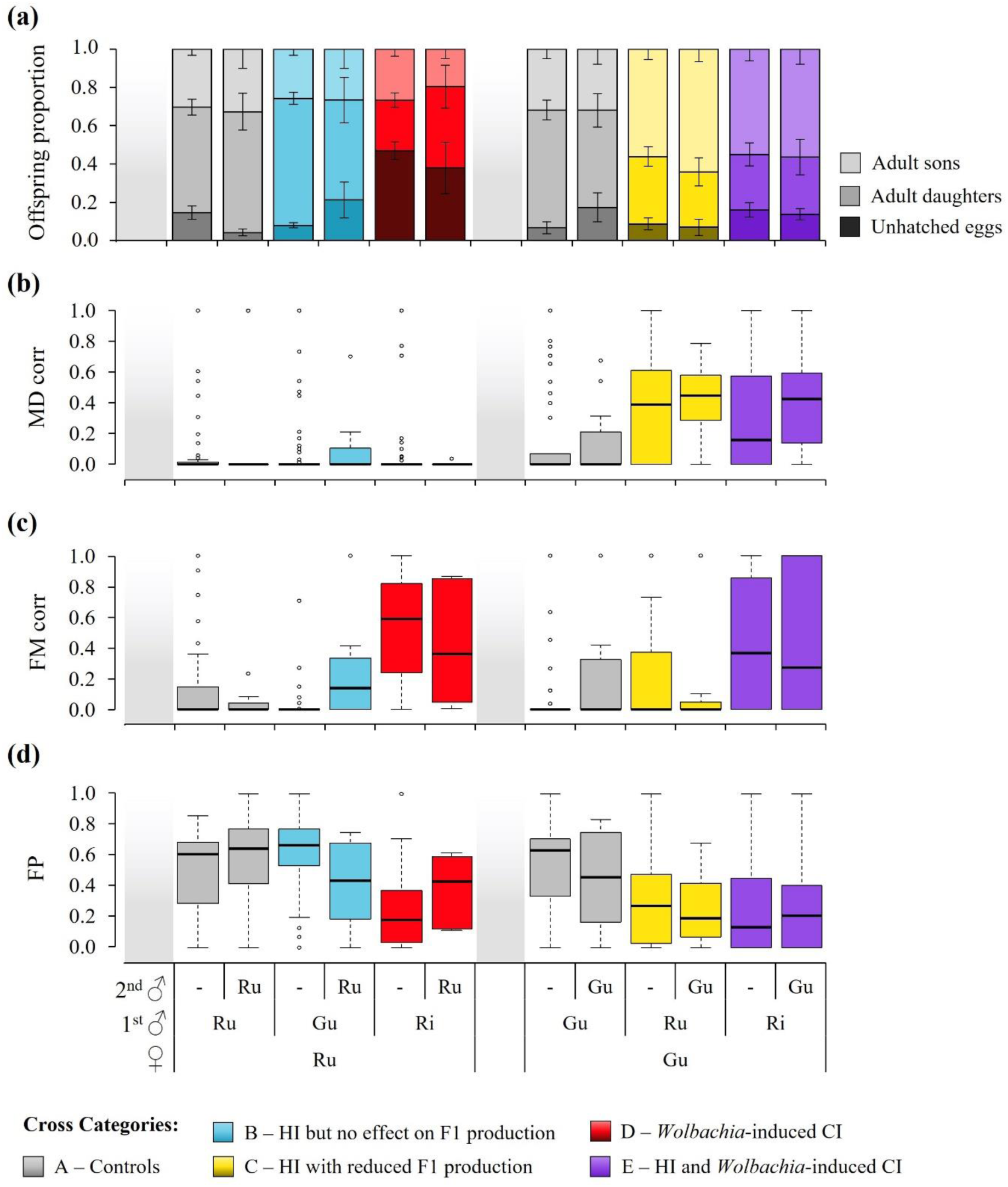
Effect of re-mating on offspring production in crosses affected by *Wolbachia*-induced CI and/or HI. (a) Outcome of egg development from each cross category, with bars representing the mean (± s.e.) relative proportions of unhatched eggs (*i.e.* embryonic mortality), adult daughters and sons. (b) Boxplot of the proportion of males produced in all crosses relative to control crosses (MD_corr_). (c) Boxplot of the proportion of estimated unhatched female eggs relative to control crosses (FM_corr_). (d) Boxplot of the proportion of F1 adult females in the brood (FP). The population of the female (♀) is displayed at the bottom level of the x-axis, that of the first male (1^st^ ♂) at the middle level, and that of the second male (2^nd^ ♂) at the top level (“-” indicates females that did not mate with the second male). CI: cytoplasmic incompatibility; HI: host-associated incompatibility.

### Contribution of intrinsic and *Wolbachia*-induced reproductive barriers to reducing gene flow

Although hybrid breakdown is the strongest reproductive barrier in both directions of crosses between the two studied spider mite populations (100% F2 embryonic mortality; Cruz et al. 2021), it ultimately contributes very little to total isolation due to the occurrence of earlier barriers (Figure 6 and Table S11). Red females and green males are mainly isolated due to hybrid sterility (98 to 100% isolation regardless of *Wolbachia* infection), as no other reproductive barrier exists in this cross direction. However, despite having the same strength in both directions of heterotypic crosses, hybrid sterility acts along with other reproductive barriers in crosses between green females and red males, which strongly reduced its contribution to total isolation (*ca.* 12% and 29% in crosses with infected and uninfected males, respectively). In this cross direction, our estimations revealed that assortative mating and fertilization failure are in fact the main sources of reproductive isolation, contributing to 27-61% and 23-71% of total isolation, respectively. Moreover, although hybrid inviability caused by the CI-inducing *Wolbachia* strain infecting red males only has a weak contribution to total isolation in heterotypic crosses as compared to homotypic crosses (*ca.* 5.5 to 6.4% in crosses between green females and Ri males *vs* 32% in crosses between Ru females and Ri males; Table S11), infection of males with this *Wolbachia* strain clearly potentiates pre-mating isolation (Figure 6). The strength of assortative mating increases from *ca.* 27% in crosses between Gu females and Ru males (non-significantly different from random mating; see Figure 1 and above) to *ca.* 61% in crosses between Gu females and Ri males (Figure 6) and to *ca.* 49% in crosses between Gi females and Ri males (Table S11).

**Figure 6.**
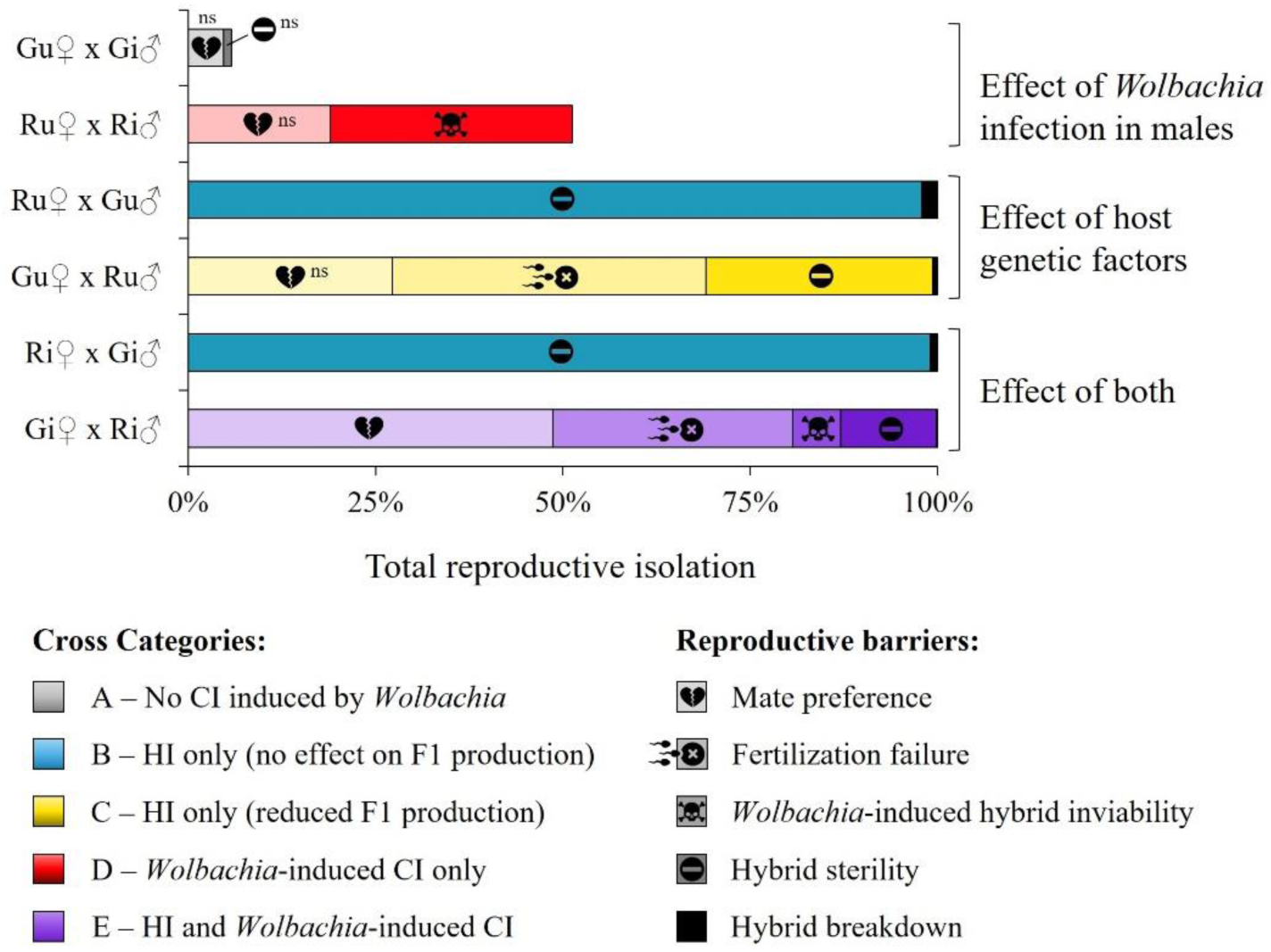
Contribution of the different reproductive barriers to reducing gene flow within and between populations. Percent contributions to total reproductive isolation were computed based on the estimated strength of reproductive isolation (*RI*) caused by a given reproductive barrier. They are shown for the six most representative types of cross in this system (see Table S11 for all crosses). ‘ns’ indicates no significant difference to zero at the 5% level. CI: cytoplasmic incompatibility; HI: host-associated incompatibility.

## Discussion

In this study, we sought to shed light on the potential role played by *Wolbachia* as an agent of pre-zygotic isolation between genetically differentiated colour forms of the spider mite *Tetranychus urticae*. To this aim, we assessed the relative contribution of *Wolbachia*-induced and host-related (pre- and post-mating) pre-zygotic barriers to total reproductive isolation. Our results revealed that *Wolbachia* infection had no effect on the mating preference of both males and females in homotypic crosses, but the CI-inducing strain infecting the red form enhanced colour-based mate preferences. Whereas both types of males showed a preference for red females, this preference seemingly disappeared when red males were cured from *Wolbachia* infection. In line with this, females showed no mate preferences in the absence of *Wolbachia* infection, but uninfected red females showed a preference for green infected males (which do not carry a CI-inducing *Wolbachia* strain) over red infected ones (which do carry a CI-inducing *Wolbachia* strain). We also found that (i) females that had engaged in matings where both types of incompatibility occurred (*Wolbachia*-induced and host-associated) were more likely to re-mate with a compatible male, and (ii) females exposed to either type of incompatibility did not significantly increase their latency to re-mate, nor reduce their copulation duration when re-mating, as compared to their first mating. Yet, we found no evidence of sperm contribution by the second (compatible) males following copulations with incompatible mates, which indicates that ‘homotypic sperm precedence’ was not a reproductive barrier at play in our experiment. Finally, our estimations of the relative contribution of each reproductive barrier to reproductive isolation between the studied populations clearly illustrate the strong asymmetries that occur in this system: red females are isolated from green males due to late intrinsic post-zygotic barriers (hybrid sterility and hybrid breakdown), whereas green females are isolated from red males by a combination of early and late reproductive barriers (pre-mating, post-mating pre-zygotic, and post-zygotic), either directly caused (hybrid inviability due to CI) or strengthened (assortative mating) by *Wolbachia* infection in red males.

### A system driven by male rather than female mate preferences

In most tested scenarios, *T. urticae* females did not choose between mates with different colour forms or infection status. This corroborates earlier results (Murtaugh and Wrensch 1978, Zhao et al. 2013b; Rodrigues et al. 2022, but see Vala et al. 2004; Rohrscheib et al. 2015), and indicates no differences in male competitive ability as well (Murtaugh and Wrensch 1978; Wagner 1998). In fact, several other studies revealed an absence of mate choice in spider mite females (*e.g.*, Magalhães et al. 2009; Zhou et al. 2020). This is surprising, as females invest more energy than males in their reproduction (Kokko et al. 2006), and spider mites have first-male sperm precedence (Helle 1967; Satoh et al. 2001; Rodrigues et al. 2020), hence the choice of the first male has enormous consequences for females (Wittenberger and Tilson 1980; Howlett 1988; Griffith et al. 2011). Possibly, this weak female choice is a consequence of male guarding of females just before their emergence as virgin adults (Potter et al. 1976), leading to little opportunity for females to choose their mate (Everson and Addicott 1982; Oku 2014). In contrast, we found strong mate preferences in males, which is also in line with earlier studies on spider mites (*e.g.*, Everson and Addicott 1982; Rodrigues et al. 2017), and in other arthropods in which males invest time and energy in pre- and/or post-copulatory guarding (reviewed in Bonduriansky 2001).

### Asymmetric reinforcement could explain the match between pre- and early post-mating barriers

In this system, one might expect assortative mating (*i.e.*, homotypic preference in both cross directions) to be selected for due to severe costs of hybridization in both cross directions (Cruz et al. 2021). Instead, our results revealed an asymmetry in pre-mating isolation (only red males prefer homotypic females). Possibly, post-mating pre-zygotic barriers (*e.g.*, fertilization failure due to cytonuclear incompatibilities, such as mitochondria-nucleus interactions; see Hill 2015) first evolved incidentally between green-form females and red-form males in allopatric populations. The resulting asymmetrical maladaptive hybridization may have subsequently (*i.e.*, at secondary contact) led to asymmetrical levels of reinforcement in areas of sympatry (Noor 1999; Servedio and Noor 2003; Coyne and Orr 2004), thereby driving the evolution of homotypic mate preferences by red males only (*e.g.*, as observed between *Drosophila recens* and *D. subquinaria* populations due to unidirectional CI induced by *Wolbachia*; Jaenike et al. 2006). This may explain the match between pre-mating and early-acting post-mating barriers in this system (sex ratio distortion likely due to fertilization failure in crosses between red males and green females; Cruz et al. 2021), as found in other systems (reviewed in Ortiz-Barrientos et al. 2009; see also Giesbers et al. 2013; Yukilevich et al. 2018). Alternatively, asymmetric barriers acting in the same cross direction could be due to genetic linkage between barriers (*e.g.*, Merrill et al. 2011), a possibility not yet investigated in spider mites. Subsequently, the two forms might have further diverged due to limited gene flow, leading to the establishment of strong late post-zygotic barriers in both directions (Servedio and Sætre 2003). In line with this, previous work has shown that barriers acting early in reproduction tend to evolve faster than those acting later (Coyne and Orr 1989; Servedio 2001; Turissini et al. 2018).

Aside from pre-mating isolation, reinforcement could also drive the evolution of other types of pre-zygotic barriers, including those occurring after mating, such as conspecific sperm precedence (Castillo and Moyle 2019; Coughlan and Matute 2020). Although preferential use of sperm from conspecific (or ‘homotypic’) males would reduce the negative effects of mating with heterospecifics (*e.g.*, Price 1997; Fricke and Arnqvist 2004; Noriyuki et al. 2012), we did not find any evidence for such reproductive barrier. Yet, green-form females previously mated with red-form males remained as receptive as when they were virgins, conversely to females mated with fully-compatible males, which became less receptive to subsequent males (increased latency and reduced copulation duration; in line with the first-male sperm precedence pattern; Helle 1967; Rodrigues et al. 2020). Similar results were also found for spider mite females first mated with (fully or partially) incompatible males of the same or different species (Clemente et al. 2016; Costa et al. 2023), but contrarily to these earlier studies, the results obtained here do not indicate any use of the sperm from second males. However, this pattern may be jeopardized under other conditions than those tested in the current study. For instance, we allowed for several copulations with the first male, and mated females were exposed to a second male only 24 hours later. This was done to detect potential issues with sperm transfer or storage when an excess of male offspring is found (*i.e.*, in crosses between green females and red males), in which case, significant effects of double mating on offspring production could not be unambiguously attributed to changes in the sperm precedence patterns (García-González 2004). However, the timing used might have been excessive to enable the use of the sperm from the second male (Potter and Wrensch 1978; Satoh et al. 2001). Moreover, given the reduced receptivity of females to second mating, the sample sizes for females mated with two different males were sometimes very low (see Table S7), which may have masked small changes in offspring production. Future studies are thus necessary to uncover potential benefits of the behaviours observed here.

### *Wolbachia*-induced CI might strengthen asymmetrical reinforcement

We show that *Wolbachia* infection strengthens assortative mating between genetically differentiated hosts, corroborating earlier findings in other systems (*e.g.*, Jaenike et al. 2006; Koukou et al. 2006; Miller et al. 2010). In addition, *Wolbachia* infection alone (*i.e.*, in homotypic crosses) has no significant effect on mate choice (as found in Rodrigues et al. 2022). Avoidance of CI might evolve more readily in structured populations, where the infection may become associated with pre-existing host traits that can be used for mate recognition (Engelstädter and Telschow 2009). Although no study has specifically addressed the population structure of spider mite populations in the field, their reliance of cultivated annual crops and several indirect lines of evidence suggest that these populations are highly structured (Navajas et al. 2000; Uesugi et al. 2009). Moreover, the fact that mating preferences contributed more to total reproductive isolation when infected red males, which carry a CI-inducing *Wolbachia* strain, were involved (see Figure 6), suggests that CI could be a mechanism driving asymmetrical reinforcement between spider-mite colour forms. Consistent with a previous study on incompatibilities between different geographic strains of green-form *T. urticae*, in which the only females receptive to a second mate were those previously mated with a genetically incompatible male carrying a CI-inducing *Wolbachia* strain (Navajas et al. 2000), we also found that only uninfected green females previously mated with a red infected male (hence carrying a CI-inducing *Wolbachia* strain) were as likely to mate with a second male as when they were virgins. In line with this, only when uninfected females (both red and green) had mated with an infected red male (with the CI-inducing strain) did their latency to copulation and copulation duration remain as when they were virgin. Together these findings thus revealed that *Wolbachia* can affect other mating behaviours beyond mating preferences (as in other systems; reviewed in Bi and Wang 2020), and raised the possibility that *Wolbachia*-induced CI could assist reinforcement processes in this system.

Whether the pattern observed in our study is only incidental or the result of *Wolbachia*-assisted reinforcement remains elusive. Indeed, while the latter hypothesis necessarily hinges upon a common *Wolbachia*-host evolutionary history, hence stable and long-lasting *Wolbachia* infection, we do not have sufficient knowledge about the past evolutionary history of the populations studied here to adjudicate which hypothesis holds true, if any. Nevertheless, previous work has shown a lack of congruence between the phylogenies of *Wolbachia* and its spider mite hosts, irrespective of their colour forms (Xie et al. 2006), or even species (Zhang et al. 2013). This suggests that *Wolbachia* infections were acquired after the forms diverged, which would preclude the bacteria from playing a role in the evolution of host reproductive barriers. In line with this, no link was found between CI induction by *Wolbachia* (neither in whether or not strains induce CI, nor in the level of CI) and the colour form of the spider mites hosting them (Gotoh et al. 2007). Moreover, the occurrence, strength, and direction of asymmetries in genetic incompatibilities between colour forms also varies depending on the host population (*e.g.*, Xue et al. 2023 vs. Cruz et al. 2021). This indicates that the pattern observed in our study is population-specific, and that *Wolbachia* is unlikely to have played a role in the establishment of incompatibilities between *T. urticae* colour forms. However, it is still possible for *Wolbachia* to be involved in the reinforcement of reproductive barriers between particular populations. Although *Wolbachia* infections can be labile in some spider mite populations, others could last much longer, depending on the transmission rate, fitness effect on host, and CI levels induced by *Wolbachia* (Zélé et al. 2020). In line with this, two subsequent field surveys conducted in the region of Lisbon, where our populations were collected, show that the prevalence of *Wolbachia* infection in red-form populations remains very high through time and across different host plant species (Zélé et al. 2018a,b), although studies on the stability of specific *Wolbachia* strains are still lacking. Alternatively, even if *Wolbachia* infections are transient, the hosts themselves may be involved in expressing the pattern of asymmetrical *Wolbachia*-induced CI observed in this study, regardless of which specific strain is infecting them. In particular, a recent study showed direct evidence that the expression of CI has a strong host-specific component in spider mites, with the host genotypes strongly modulating the expression (level and pattern) of CI induced by a same *Wolbachia* strain (Wybouw et al. 2022). This should be of particular relevance for host populations living in areas where *Wolbachia* infections are highly prevalent, as in the case of spider mites from the Iberian Peninsula (Zélé et al. 2018a,b; Pina et al. 2020).

### Not just a missing barrier: Heterotypic mate preference may be an adaptive strategy

Although reinforcement is a seductive hypothesis to explain why red-form males prefer red females, it does not explain why green-form males also prefer these females. The occurrence of such seemingly maladaptive behaviour suggests that other, or additional, evolutionary forces are at play.

One possibility could be that heterotypic mating preference is a by-product of inbreeding avoidance in the green-form population. Spider mites effectively avoid related individuals (Tien et al. 2011; Bitume et al. 2013; Yoshioka and Yano 2014), but it is not clear whether this extends to more distantly-related individuals. For instance, males of both *T. evansi* and *T. urticae* preferentially mate with *T. urticae* females (Sato et al. 2016; but see Clemente et al. 2016), but this occurs even when *T. evansi* females are non-kin (Sato et al. 2016). Moreover, this supposes that the green population suffers more from inbreeding than the red one, a possibility that could be tested in the future.

Another possibility could be that preference of both types of males for red-form females is due to these females being more attractive. For instance, a new trait (*e.g.*, a pheromone profile) may have evolved in red females in response to intense female competition (*i.e.*, their sex ratio is more female-biased than that of green mites when they oviposit in groups; unpublished data), and this trait may then be fortuitously preferred by green males if it stimulates the same coding system as the ancestral trait (Endler and Basolo 1998). Alternatively, both types of male may have conserved an ancestral preference for a trait that has been lost or diverged in green-form females (Endler and Basolo 1998). This could occur if the rate of evolution of male preference is slower than that of the female trait. The observed male preferences may also be caused by differences in female reluctance and male vigour (*e.g.*, van den Berg et al. 1984) in response to stronger sexual conflicts in the green-form population. This hypothesis is supported by the fact that green females are less likely to mate than red females even in the absence of choice, whereas green males spend longer periods of time copulating than red males do (suggesting longer post-copulatory guarding; (Satoh et al. 2001). In line with this, theory predicts that sexual conflicts can drive the evolution of mate preferences, increasing reproductive isolation and, consequently, the rates of speciation (Parker and Partridge 1998).

Finally, building upon the recent idea that partial reproductive isolation may be an adaptive optimum (Servedio and Hermisson 2020), we considered the possibility that heterotypic mating preference might be selected for under reproductive interference (Gröning and Hochkirch 2008), as the two colour forms have overlapping distribution and host plant range (Migeon and Dorkeld 2023), and often co-occur on the same individual host plant (Lu et al. 2017, 2018; Zélé et al. 2018b). Although most conditions that have been theoretically considered to promote the evolution of ‘disassortative mating’, such as a heterozygote advantage (*e.g.*, Maisonneuve et al. 2021), are not met in our system (hybrids are sterile or suffer breakdown; Cruz et al. 2021), heterotypic mating preference may still confer higher benefits than costs to the green-form population in the presence of red-form competitors. Indeed, this behaviour should be highly costly for red females due to first male sperm precedence, but green males may only pay relatively small costs as they can mate multiple times (Krainacker and Carey 1989). Thus, similarly to how CI induced by *Wolbachia* increases the relative fitness of infected females, the ‘spiteful’ behaviour of green males might be selected for as it confers an indirect fitness advantage to their green sisters (Hamilton 1970; Gardner and West 2004; Engelstädter and Charlat 2006). Disassortative mating may thus act synergistically with sex-ratio distortion (*i.e.*, the overproduction of sons) in crosses between green females and red males (see Cruz et al. 2021) to promote the exclusion of the red form population (see Grether et al. 2017; Cruz et al. 2023). Conversely, homotypic mating preference by red males should decrease the strength of reproductive interference for the red population, as it reduces the prevalence of crosses between green females and red males (hence the overproduction of green males stemming from these crosses) and should prevent (non-choosy) red females from having a higher chance to mate with a green male. Following this hypothesis, the CI-inducing *Wolbachia* strain naturally infecting the red-form population seems to favour its own host population by increasing the likelihood that red males mate with compatible (red) females, whereas it has no control over heterotypic mating preference by green males. Testing whether such an ‘escalating arms race’ could indeed occur in response to reproductive interference (involving or not *Wolbachia*-induced CI) is of high relevance for future speciation studies.

## Conclusion

In this study, we identified a mechanism through which *Wolbachia* could assist host speciation processes. Our results show that *Wolbachia* infection in *T. urticae* males indirectly contributes to pre-mating isolation between genetically differentiated *T. urticae* colour forms by strengthening pre-existing preferences. These preferences match early post-mating barriers in the system, as crosses that are affected both by host-associated and *Wolbachia*-induced incompatibilities are generally avoided. Our results also further highlight the importance of pre-mating isolation in this system, as they revealed that, in our experimental conditions, females of either form are unable to compensate for incompatible crosses by re-mating. Overall, our comprehensive study of pre- and post-zygotic reproductive barriers allowed identifying asymmetries in patterns of isolation between the two populations, hinting at a possible history of reinforcement followed by an interruption of gene flow. These findings also open new research avenues, such as to study the impact of complex patterns of isolation on population dynamics, and of the resulting selection pressures on the evolutionary trajectories of populations.

## Authors’ contributions

MC, SM, and FZ conceived and designed the experiments. MC and MB performed the choice and no-choice tests, respectively. MC and FZ analysed the data. Funding agencies did not participate in the design or analysis of experiments. MC and FZ wrote the manuscript with input from SM. All authors read and approved the final version of the manuscript.

## Acknowledgements

We are grateful to Inês Santos for the maintenance of the spider mite populations and the plants, to Leonor Rodrigues and Élio Sucena for useful advises with experimental designs, and to Vitor Sousa and Alexandre Blanckaert for useful comments on an early version of the manuscript. We would like to especially thank the Associate Editor Dr. Brandon S. Cooper, as well as two anonymous reviewers, for their very constructive and detailed comments, which helped us greatly improve the manuscript.

## Funding

This work was funded by an FCT-ANR project (FCT-ANR//BIA-EVF/0013/2012) to SM and Isabelle Olivieri, and by an ERC Consolidator Grant (COMPCON, GA 725419) to SM. MC was funded through an FCT PhD fellowship (SFRH/BD/136454/2018), and FZ through an FCT Post-Doc fellowship (SFRH/BPD/125020/2016) when experiments were performed. This is contribution ISEM-2024-XXX of the Institute of Evolutionary Science of Montpellier (ISEM). For the purposes of Open Access, a CC-BY 4.0 public copyright licence has been applied by the authors to the present document and will be applied to all subsequent versions up to the Author Accepted Manuscript arising from this submission.

## Conflict of interest

The authors declare that they have no conflict of interest with the content of this article.

## Data availability

All datasets and R scripts are available at Zenodo (https://doi.org/10.5281/zenodo.11160702)

## Notes

https://doi.org/10.5281/zenodo.11160702

### Competing Interest Statement

The authors have declared no competing interest.

### Summary of Updates

Several minor changes were made, especially in the Introduction/Discussion, to improve the clarity and quality of the manuscript.

## References

Auger, P., A. Migeon, E. A. Ueckermann, L. Tiedt, and M. Navajas. 2013. Evidence for synonymy between *Tetranychus urticae* and *Tetranychus cinnabarinus* (Acari, Prostigmata, Tetranychidae): Review and new data. Acarologia 53:383–415.

Baack, E., M. C. Melo, L. H. Rieseberg, and D. Ortiz-Barrientos. 2015. The origins of reproductive isolation in plants. New Phytol 207:968–984.

Bank, C., J. Hermisson, and M. Kirkpatrick. 2012. Can reinforcement complete speciation? Evolution 66:229–239.

Bateman, A. J. 1949. Analysis of data on sexual isolation. Evolution 3:174–177.

Bhat, S., P. A. Amundsen, R. Knudsen, K. Ø. Gjelland, S. E. Fevolden, L. Bernatchez, and K. Præbel. 2014. Speciation reversal in European whitefish (*Coregonus lavaretus* (L.)) caused by competitor invasion. PLoS One 9:e91208.

Bi, J., and Y. F. Wang. 2020. The effect of the endosymbiont *Wolbachia* on the behavior of insect hosts. Insect Sci 27:846–858.

Bitume, E. V., D. Bonte, O. Ronce, F. Bach, E. Flaven, I. Olivieri, and C. M. Nieberding. 2013. Density and genetic relatedness increase dispersal distance in a subsocial organism. Ecol Lett 16:430–437.

Bonduriansky, R. 2001. The evolution of male mate choice in insects: a synthesis of ideas and evidence. Biol Rev 76:305–339.

Bordenstein, S., F. P. O’Hara, and J. H. Werren. 2001. *Wolbachia*-induced incompatibility precedes other hybrid incompatibilities in *Nasonia*. Nature 409:707–710.

Breeuwer, J. A. J. 1997. *Wolbachia* and cytoplasmic incompatibility in the spider mites *Tetranychus urticae* and *T. turkestani*. Heredity 79:41–47.

Breeuwer, J. A. J., and G. Jacobs. 1996. *Wolbachia*: Intracellular manipulators of mite reproduction. Exp Appl Acarol 20:421–434.

Breeuwer, J. A. J., and J. H. Werren. 1990. Microorganisms associated with chromosome destruction and reproductive isolation between two insect species. Nature 346:558–560.

Brucker, R. M., and S. R. Bordenstein. 2012. Speciation by symbiosis. Trends Ecol Evol 27:443–451.

Bruzzese, D. J., H. Schuler, T. M. Wolfe, M. M. Glover, J. Mastroni, M. M. Doellman, C. Tait, W. L. Yee, J. Rull, M. Aluja, G. R. Hood, R. Goughnour, C. Stauffer, P. Nosil, and J. L. Feder. 2022. Testing the potential contribution of *Wolbachia* to speciation when cytoplasmic incompatibility becomes associated with host-related reproductive isolation. Mol Ecol 31:2935–2950.

Castillo, D. M., and L. C. Moyle. 2019. Conspecific sperm precedence is reinforced, but postcopulatory sexual selection weakened, in sympatric populations of *Drosophila*. Proc R Soc B 286:20182535.

Chain-Ing, T. S., and S. Sheuan-Ping. 1995. Copulation competition of male *Tetranychus urticae* Koch and *Tetranychus kanzawai* Kishida (Acarina: Tetranychidae) with conspecific and heterospecific females and their isolation mechanism. Chin J Entomol 15:239–255.

Champion de Crespigny, F. E., R. K. Butlin, and N. Wedell. 2005. Can cytoplasmic incompatibility inducing *Wolbachia* promote the evolution of mate preferences? J Evol Biol 18:967–977.

Champion de Crespigny, F. E., and N. Wedell. 2006. *Wolbachia* infection reduces sperm competitive ability in an insect. Proc R Soc B 273:1455–1458.

Champion de Crespigny, F. E., L. D. Hurst, and N. Wedell. 2008. Do *Wolbachia*-associated incompatibilities promote polyandry? Evolution 62:107–122.

Clemente, S. H., L. R. Rodrigues, R. Ponce, S. A. M. Varela, and S. Magalhães. 2016. Incomplete species recognition entails few costs in spider mites, despite first-male precedence. Behav Ecol Sociobiol 70:1161–1170.

Clemente, S. H., I. Santos, R. Ponce, L. R. Rodrigues, S. A. M. Varela, and S. Magalhães. 2018. Despite reproductive interference, the net outcome of reproductive interactions among spider mite species is not necessarily costly. Behav Ecol 29:321–327.

Cooper, B. S., P. S. Ginsberg, M. Turelli, and D. R. Matute. 2017. *Wolbachia* in the *Drosophila yakuba* complex: Pervasive frequency variation and weak cytoplasmic incompatibility, but no apparent effect on reproductive isolation. Genetics 205:333–351.

Costa, S. G., S. Magalhães, and L. R. Rodrigues. 2023. Multiple mating rescues offspring sex ratio but not productivity in a haplodiploid exposed to developmental heat stress. Funct Ecol 37:1291–1303.

Coughlan, J. M., and D. R. Matute. 2020. The importance of intrinsic postzygotic barriers throughout the speciation process. Phil Trans R Soc B 375:20190533.

Coyne, J. A., and H. A. Orr. 1989. Patterns of speciation in *Drosophila*. Evolution 43:362–381.

Coyne, J. A., and H. A. Orr. 1997. “Patterns of speciation in Drosophila” revisited. Evolution 51:295–303.

Coyne, J. A., and H. A. Orr. 2004. Speciation. Sinauer, Sunderland, MA.

Crawley, M. J. 2007. The R Book. John Wiley & Sons, Ltd, Chichester.

Cruz, M. A., O. Godoy, I. Fragata, V. C. Sousa, S. Magalhães, and F. Zélé. 2023. Sex in the kitchen: non-additive effects of competition for food and reproductive interference on coexistence outcomes between closely related species. doi: 10.1101/2023.11.09.566372.

Cruz, M. A., S. Magalhães, É. Sucena, and F. Zélé. 2021. *Wolbachia* and host intrinsic reproductive barriers contribute additively to postmating isolation in spider mites. Evolution 75:2085–2101.

de Boer, R. 1982a. Laboratory hybridization between semi-incompatible races of the arrhenotokous spider mite Tetranychus urticae Koch (Acari: Tetranychidae). Evolution 36:553–560.

de Boer, R. 1982b. Partial hybrid sterility between strains of the arrhenotokous spider mite, Tetranychus urticae complex (Acari, Tetranychidae). Genetica 58:23–33.

Dupont, L. M. 1979. On gene flow between *Tetranychus urticae* Koch, 1836 and *Tetranychus cinnabarinus* (Boisduval) Boudreaux, 1956 (Acari: Tetranychidae): Synonymy between the two species. Entomol Exp Appl 25:297–303.

Duron, O., D. Bouchon, S. S. S. Boutin, L. Bellamy, L. Zhou, J. Engelstädter, and G. D. Hurst. 2008. The diversity of reproductive parasites among arthropods: *Wolbachia* do not walk alone. BMC Biol 6:27.

Endler, J., and A. Basolo. 1998. Sensory ecology, receiver biases and sexual selection. Trends Ecol Evol 13:415–420.

Engelstädter, J., and S. Charlat. 2006. Outbreeding selects for spiteful cytoplasmic elements. Proc R Soc B 273:923–929.

Engelstädter, J., and G. D. D. Hurst. 2009. The ecology and evolution of microbes that manipulate host reproduction. Annu Rev Ecol Evol Syst 40:127–149.

Engelstädter, J., and A. Telschow. 2009. Cytoplasmic incompatibility and host population structure. Heredity 103:196–207.

Everson, P. R., and J. F. Addicott. 1982. Mate selection strategies by male mites in the absence of intersexual selection by females: a test of six hypotheses. Can J Zool 60:2729–2736.

Fortin, M., C. Debenest, C. Souty-Grosset, and F. J. Richard. 2018. Males prefer virgin females, even if parasitized, in the terrestrial isopod *Armadillidium vulgare*. Ecol Evol 8:3341–3353.

Fricke, C., and G. Arnqvist. 2004. Conspecific sperm precedence in flour beetles. Anim Behav 67:729–732.

García-González, F. 2004. Infertile matings and sperm competition: the effect of “nonsperm representation” on intraspecific variation in sperm precedence patterns. Am Nat 164:457–472.

Gardner, A., and S. A. West. 2004. Spite and the scale of competition. J Evol Biol 17:1195– 1203.

Giesbers, M. C. W. G., S. Gerritsma, J. Buellesbach, W. Diao, B. A. Pannebakker, L. van de Zande, T. Schmitt, and L. W. Beukeboom. 2013. Prezygotic isolation in the parasitoid wasp genus *Nasonia*. Pp. 165–192 *in* P. Michalak, ed. Speciation: Natural Processes, Genetics and Biodiversity. Nova Biomedical, New York.

Godinho, D. P., M. A. Cruz, M. Charlery de la Masselière, J. Teodoro-Paulo, C. Eira, I. Fragata, L. R. Rodrigues, F. Zélé, and S. Magalhães. 2020. Creating outbred and inbred populations in haplodiploids to measure adaptive responses in the laboratory. Ecol Evol 10:7291–7305.

Gotoh, T., J. Bruin, M. W. Sabelis, and S. B. J. Menken. 1993. Host race formation in *Tetranychus urticae*: genetic differentiation, host plant preference, and mate choice in a tomato and a cucumber strain. Entomol Exp Appl 68:171–178.

Gotoh, T., H. Noda, and X. Y. Hong. 2003. *Wolbachia* distribution and cytoplasmic incompatibility based on a survey of 42 spider mite species (Acari: Tetranychidae) in Japan. Heredity 91:208–216.

Gotoh, T., J. Sugasawa, H. Noda, and Y. Kitashima. 2007. *Wolbachia*-induced cytoplasmic incompatibility in Japanese populations of *Tetranychus urticae* (Acari: Tetranychidae). Exp Appl Acarol 42:1–16.

Grether, G. F., K. S. Peiman, J. A. Tobias, and B. W. Robinson. 2017. Causes and consequences of behavioral interference between species. Trends Ecol Evol 32:760–772.

Griffith, S. C., S. R. Pryke, and W. A. Buttemer. 2011. Constrained mate choice in social monogamy and the stress of having an unattractive partner. Proc R Soc B 278:2798– 2805.

Gröning, J., and A. Hochkirch. 2008. Reproductive interference between animal species. Q Rev Biol 83:257–282.

Hamilton, W. D. 1970. Selfish and spiteful behaviour in an evolutionary model. Nature 228:1218–1220.

He, Z., H. B. Zhang, S. T. Li, W. J. Yu, J. Biwot, X. Q. Yu, Y. Peng, and Y. F. Wang. 2018. Effects of *Wolbachia* infection on the postmating response in *Drosophila melanogaster*. Behav Ecol Sociobiol 72:146.

Helle, W. 1967. Fertilization in the two-spotted spider mite (*Tetranychus urticae*: Acari). Entomol Exp Appl 10:103–110.

Helle, W., and H. R. Bolland. 1967. Karyotypes and sex-determination in spider mites (Tetranychidae). Genetica 38:43–53.

Helle, W., and C. F. Van de Bund. 1962. Crossbreeding experiments with some species of the *Tetranychus urticae* group. Entomol Exp Appl 5:159–165.

Hill, G. E. 2015. Mitonuclear ecology. Mol Biol Evol 32:1917–1927.

Hill, R. L., and D. J. O’Donnell. 1991. Reproductive isolation between *Tetranychus lintearius* and two related mites, T. urticae and T. turkestani (Acarina: Tetranychidae). Exp Appl Acarol 11:241–251.

Hinomoto, N., M. Osakabe, T. Gotoh, and A. Takafuji. 2001. Phylogenetic analysis of green and red forms of the two-spotted spider mite, *Tetranychus urticae* Koch (Acari: Tetranychidae), in Japan, based on mitochondrial cytochrome oxidase subunit I sequences. Appl Entomol Zool 36:459–464.

Howlett, R. 1988. Sexual selection by female choice in monogamous birds. Nature 332:583– 584.

Jaenike, J., K. A. Dyer, C. Cornish, and M. S. Minhas. 2006. Asymmetrical reinforcement and *Wolbachia* infection in *Drosophila*. PLoS Biol 4:1852–1862.

Kaur, R., J. D. Shropshire, K. L. Cross, B. Leigh, A. J. Mansueto, V. Stewart, S. R. Bordenstein, and S. R. Bordenstein. 2021. Living in the endosymbiotic world of *Wolbachia*: a centennial review. Cell Host Microbe 29:879–893.

Kearns, A. M., M. Restani, I. Szabo, A. Schrøder-Nielsen, J. A. Kim, H. M. Richardson, J. M. Marzluff, R. C. Fleischer, A. Johnsen, and K. E. Omland. 2018. Genomic evidence of speciation reversal in ravens. Nat Commun 9:906.

Keh, B. 1952. Mating experiments with the two-spotted spider mite complex. J Econ Entomol 45:308–312.

Knegt, B., T. Potter, N. A. Pearson, Y. Sato, H. Staudacher, B. C. J. Schimmel, E. T. Kiers, and M. Egas. 2017. Detection of genetic incompatibilities in non-model systems using simple genetic markers: hybrid breakdown in the haplodiploid spider mite *Tetranychus evansi*. Heredity 118:311–321.

Kokko, H., M. D. Jennions, and R. Brooks. 2006. Unifying and testing models of sexual selection. Annu Rev Ecol Evol Syst 37:43–66.

Koukou, K., H. Pavlikaki, G. Kilias, J. H. Werren, K. Bourtzis, and S. N. Alahiotis. 2006. Influence of antibiotic treatment and *Wolbachia* curing on sexual isolationg among *Drosophila melanogaster* cage populations. Evolution 60:87–96.

Krainacker, D. A., and J. R. Carey. 1989. Reproductive limits and heterogeneity of male twospotted spider mites. Entomol Exp Appl 50:209–214.

Lackey, A. C. R., and J. W. Boughman. 2017. Evolution of reproductive isolation in stickleback fish. Evolution 71:357–372.

Lewis, Z., F. E. C. De Crespigny, S. M. Sait, T. Tregenza, and N. Wedell. 2011. *Wolbachia* infection lowers fertile sperm transfer in a moth. Biol Lett 7:187–189.

Liu, C., J. L. Wang, Y. Zheng, E. J. Xiong, J. J. Li, L. L. Yuan, X. Q. Yu, and Y. F. Wang. 2014. *Wolbachia*-induced paternal defect in *Drosophila* is likely by interaction with the juvenile hormone pathway. Insect Biochem Mol Biol 49:49–58.

Lu, W., Y. Hu, P. Wei, Q. Xu, C. Bowman, M. Li, and L. He. 2018. Acaricide-mediated competition between the sibling species *Tetranychus cinnabarinus* and *Tetranychus urticae*. J Econ Entomol 111:1346–1353.

Lu, W., M. Wang, Z. Xu, G. Shen, P. Wei, M. Li, W. Reid, and L. He. 2017. Adaptation of acaricide stress facilitates *Tetranychus urticae* expanding against *Tetranychus cinnabarinus* in China. Ecol Evol 7:1233–1249.

Magalhães, S., E. Blanchet, M. Egas, and I. Olivieri. 2009. Are adaptation costs necessary to build up a local adaptation pattern? BMC Evol Biol 9:182.

Maisonneuve, L., T. Beneteau, M. Joron, C. Smadi, and V. Llaurens. 2021. When do opposites attract? A model uncovering the evolution of disassortative mating. Am Nat 198:625– 641.

Malogolowkin-Cohen, Ch., A. S. Simmons, and H. Levene. 1965. A study of sexual isolation between certain strains of *Drosophila paulistorum*. Evolution 19:95–103.

Matsuda, T., T. Kozaki, K. Ishii, and T. Gotoh. 2018. Phylogeny of the spider mite sub-family Tetranychinae (Acari: Tetranychidae) inferred from RNA-Seq data. PLoS One 13:e0203136.

Mayr, E. 1942. Systematics and the Origin of Species. Columbia University Press, New York.

Merrell, D. J. 1950. Measurement of Sexual Isolation and Selective Mating. Evolution 4:326–331.

Merrill, R. M., B. Van Schooten, J. A. Scott, and C. D. Jiggins. 2011. Pervasive genetic associations between traits causing reproductive isolation in *Heliconius butterflies*. Proc R Soc B 278:511–518.

Migeon, A., and F. Dorkeld. 2023. Spider Mites Web: a comprehensive database for the Tetranychidae.

Miller, W. J., L. Ehrman, and D. Schneider. 2010. Infectious speciation revisited: impact of symbiont-depletion on female fitness and mating behavior of *Drosophila paulistorum*. PLoS Pathog 6:e1001214.

Murtaugh, M. P., and D. L. Wrensch. 1978. Interspecific competition and hybridization between twospotted and carmine spider mites. Ann Entomol Soc Am 71:862–864.

Navajas, M., A. Tsagkarakou, J. Lagnel, and M. J. Perrot-Minnot. 2000. Genetic differentiation in *Tetranychus urticae* (Acari: Tetranychidae): polymorphism, host races or sibling species? Exp Appl Acarol 24:365–376.

Noor, M. A. F. 1999. Reinforcement and other consequences of sympatry. Heredity 83:503– 508.

Noriyuki, S., N. Osawa, and T. Nishida. 2012. Asymmetric reproductive interference between specialist and generalist predatory ladybirds. J Anim Ecol 81:1077–1085.

Oku, K. 2014. Sexual selection and mating behavior in spider mites of the genus *Tetranychus* (Acari: Tetranychidae). Appl Entomol Zool 49:1–9.

Ortiz-Barrientos, D., A. Grealy, and P. Nosil. 2009. The genetics and ecology of reinforcement: implications for the evolution of prezygotic isolation in sympatry and beyond. Ann NY Acad Sci 1168:156–182

Parker, G. A., and L. Partridge. 1998. Sexual conflict and speciation. Philos Trans R Soc Lond B 353:261–274.

Perrot-Minnot, M. J., B. Cheval, A. Migeon, and M. Navajas. 2002. Contrasting effects of *Wolbachia* on cytoplasmic incompatibility and fecundity in the haplodiploid mite *Tetranychus urticae*. J Evol Biol 15:808–817.

Pina, T., B. Sabater-Muñoz, M. Cabedo-López, J. Cruz-Miralles, J. A. Jaques, and M. A. Hurtado-Ruiz. 2020. Molecular characterization of *Cardinium*, *Rickettsia*, *Spiroplasma* and *Wolbachia* in mite species from citrus orchards. Exp Appl Acarol 81:335–355.

Potter, D. A., and D. L. Wrensch. 1978. Interrupted matings and the effectiveness of second inseminations in the twospotted spider mite. Ann Entomol Soc Am 71:882–885.

Potter, D. A., D. L. Wrensch, and D. E. Johnston. 1976. Aggression and mating success in male spider mites. Science 193:160–161.

Price, C. S. C. 1997. Conspecific sperm precedence in *Drosophila*. Nature 388:663–666.

Ramsey, J., H. D. Bradshaw, and D. W. Schemske. 2003. Components of reproductive isolation between the monkeyflowers *Mimulus lewisii* and *M. cardinalis* (Phrymaceae). Evolution 57:1520–1534.

Richard, F. J. 2017. Symbiotic bacteria influence the odor and mating preference of their hosts. Front Ecol Evol 5:143.

Rodrigues, L. R., A. R. T. Figueiredo, T. Van Leeuwen, I. Olivieri, and S. Magalhães. 2020. Costs and benefits of multiple mating in a species with first-male sperm precedence. J Anim Ecol 89:1045–1054.

Rodrigues, L. R., A. R. T. Figueiredo, S. A. M. Varela, I. Olivieri, and S. Magalhães. 2017. Male spider mites use chemical cues, but not the female mating interval, to choose between mates. Exp Appl Acarol 71:1–13.

Rodrigues, L. R., F. Zélé, I. Santos, and S. Magalhães. 2022. No evidence for the evolution of mating behavior in spider mites due to *Wolbachia*-induced cytoplasmic incompatibility. Evolution 76:623–635.

Sato, Y., and J. M. Alba. 2020. Reproductive interference and sensitivity to female pheromones in males and females of two herbivorous mite species. Exp Appl Acarol 81:59–74.

Sato, Y., J. M. Alba, and M. W. Sabelis. 2014. Testing for reproductive interference in the population dynamics of two congeneric species of herbivorous mites. Heredity 113:495– 502.

Sato, Y., H. Staudacher, and M. W. Sabelis. 2016. Why do males choose heterospecific females in the red spider mite? Exp Appl Acarol 68:21–31.

Satoh, Y., S. Yano, and A. Takafuji. 2001. Mating strategy of spider mite, *Tetranychus urticae* (Acari: Tetranychidae) males: postcopulatory guarding to assure paternity. Appl Entomol Zool 36:41–45.

Schneider, D. I., L. Ehrman, T. Engl, M. Kaltenpoth, A. Hua-Van, A. Le Rouzic, and W. J. Miller. 2019. Symbiont-driven male mating success in the neotropical *Drosophila paulistorum* superspecies. Behav Genet 49:83–98.

Servedio, M. R. 2001. Beyond reinforcement: the evolution of premating isolation by direct selection on preferences and postmating, prezygotic incompatibilities. Evolution 55:1909–1920.

Servedio, M. R., and J. Hermisson. 2020. The evolution of partial reproductive isolation as an adaptive optimum. Evolution 74:4–14.

Servedio, M. R., and M. Noor. 2003. The role of reinforcement in speciation: theory and data. Annu Rev Ecol Evol Syst 34:339–364.

Servedio, M. R., and G.-P. Sætre. 2003. Speciation as a positive feedback loop between postzygotic and prezygotic barriers to gene flow. Proc R Soc Lond B 270:1473–1479.

Shoemaker, D. D., V. Katju, and J. Jaenike. 1999. *Wolbachia* and the evolution of reproductive isolation between *Drosophila recens* and *Drosophila subquinaria*. Evolution 53:1157– 1164.

Shropshire, J. D., and S. R. Bordenstein. 2016. Speciation by symbiosis: the microbiome and behavior. mBio 7:e01785–15.

Shropshire, J. D., B. Leigh, and S. R. Bordenstein. 2020. Symbiont-mediated cytoplasmic incompatibility: what have we learned in 50 years? eLife 9:e61989.

Snook, R. R., S. Y. Cleland, M. F. Wolfner, and T. L. Karr. 2000. Offsetting effects of *Wolbachia* infection and heat shock on sperm production in *Drosophila simulans*: analyses of fecundity, fertility and accessory gland proteins. Genetics 155:167–178.

Stankowski, S., and M. Ravinet. 2021. Defining the speciation continuum. Evolution 75:1256– 1273.

Sugasawa, J., Y. Kitashima, and T. Gotoh. 2002. Hybrid affinities between the green and the red forms of the two-spotted spider mite *Tetranychus urticae* (Acari: Tetranychidae) under laboratory and seminatural conditions. Appl Entomol Zool 37:127–139.

Suh, E., C. Sim, J. J. Park, and K. Cho. 2015. Inter-population variation for *Wolbachia* induced reproductive incompatibility in the haplodiploid mite *Tetranychus urticae*. Exp Appl Acarol 65:55–71.

Taylor, E. B., J. W. Boughman, M. Groenenboom, M. Sniatynski, D. Schluter, and J. L. Gow. 2006. Speciation in reverse: morphological and genetic evidence of the collapse of a three-spined stickleback (*Gasterosteus aculeatus*) species pair. Mol Ecol 15:343–355.

Telschow, A., P. Hammerstein, and J. H. Werren. 2005. The effect of *Wolbachia* versus genetic incompatibilities on reinforcement and speciation. Evolution 59:1607–1619.

Tien, N. S. H., G. Massourakis, M. W. Sabelis, and M. Egas. 2011. Mate choice promotes inbreeding avoidance in the two-spotted spider mite. Exp Appl Acarol 54:119–124.

Turissini, D. A., J. A. McGirr, S. S. Patel, J. R. David, and D. R. Matute. 2018. The rate of evolution of postmating-prezygotic reproductive isolation in *Drosophila*. Mol Biol Evol 35:312–334.

Uesugi, R., Y. Kunimoto, and M. Osakabe. 2009. The fine-scale genetic structure of the two-spotted spider mite in a commercial greenhouse. Exp Appl Acarol 47:99–109.

Vala, F., M. Egas, J. A. J. Breeuwer, and M. W. Sabelis. 2004. *Wolbachia* affects oviposition and mating behaviour of its spider mite host. J Evol Biol 17:692–700.

Vala, F., A. Weeks, D. Claessen, J. A. J. Breeuwer, and M. W. Sabelis. 2002. Within- and between-population variation for *Wolbachia*-induced reproductive incompatibility in a haplodiploid mite. Evolution 56:1331–1339.

Van de Bund, C. F., and W. Helle. 1960. Investigations on the *Tetranychus urticae* complex in north west Europe (Acari: Tetranychidae). Entomol Exp Appl 3:142–156.

van den Berg, M. J., G. Thomas, H. Hendriks, and W. van Delden. 1984. A reexamination of the negative assortative mating phenomenon and its underlying mechanism in *Drosophila melanogaster*. Behav Genet 14:45–61.

Villacis-Perez, E., S. Snoeck, A. H. Kurlovs, R. M. Clark, J. A. J. Breeuwer, and T. Van Leeuwen. 2021. Adaptive divergence and post-zygotic barriers to gene flow between sympatric populations of a herbivorous mite. Commun Biol 4:853.

Weinert, L. A., E. V. Araujo-Jnr, M. Z. Ahmed, and J. J. Welch. 2015. The incidence of bacterial endosymbionts in terrestrial arthropods. Proc R Soc B 282:20150249.

Werren, J. H., L. Baldo, and M. E. Clark. 2008. *Wolbachia*: master manipulators of invertebrate biology. Nat Rev Microbiol 6:741–751.

Wittenberger, J. F., and R. L. Tilson. 1980. The evolution of monogamy: hypotheses and evidence. Annu Rev Ecol Syst 11:197–232.

Wybouw, N., F. Mortier, and D. Bonte. 2022. Interacting host modifier systems control *Wolbachia*-induced cytoplasmic incompatibility in a haplodiploid mite. Evol Lett 6:255–265.

Xie, L., H. Miao, and X.-Y. Hong. 2006. The two-spotted spider mite *Tetranychus urticae* Koch and the carmine spider mite *Tetranychus cinnabarinus* (Boisduval) in China mixed in their *Wolbachia* phylogenetic tree. Zootaxa 1165:33–46.

Xue, W., J. Sun, J. Witters, M. Vandenhole, W. Dermauw, S. A. Bajda, E. A. Simma, N. Wybouw, E. Villacis-Perez, and T. Van Leeuwen. 2023. Incomplete reproductive barriers and genomic differentiation impact the spread of resistance mutations between green- and red-colour morphs of a cosmopolitan mite pest. Mol Ecol 32:4278–4297.

Yoshioka, T., and S. Yano. 2014. Do *Tetranychus urticae* males avoid mating with familiar females? J Exp Biol 217:2297–2300.

Yukilevich, R., L. S. Maroja, K. Nguyen, S. Hussain, and P. Kumaran. 2018. Rapid sexual and genomic isolation in sympatric *Drosophila* without reproductive character displacement. Ecol Evol 8:2852–2867.

Zélé, F., I. Santos, M. Matos, M. Weill, F. Vavre, and S. Magalhães. 2020. Endosymbiont diversity in natural populations of *Tetranychus* mites is rapidly lost under laboratory conditions. Heredity 124:603–617.

Zélé, F., I. Santos, I. Olivieri, M. Weill, O. Duron, and S. Magalhães. 2018a. Endosymbiont diversity and prevalence in herbivorous spider mite populations in South-Western Europe. FEMS Microbiol Ecol 94:fiy015.

Zélé, F., J. L. Santos, D. P. Godinho, and S. Magalhães. 2018b. *Wolbachia* both aids and hampers the performance of spider mites on different host plants. FEMS Microbiol Ecol 94: fiy187.

Zhang, Y.-K., Y.-T. Chen, K. Yang, G.-X. Qiao, and X.-Y. Hong. 2016. Screening of spider mites (Acari: Tetranychidae) for reproductive endosymbionts reveals links between co-infection and evolutionary history. Sci Rep 6:27900.

Zhang, Y.-K., K.-J. Zhang, J.-T. Sun, X.-M. Yang, C. Ge, and X.-Y. Hong. 2013. Diversity of *Wolbachia* in natural populations of spider mites (genus *Tetranychus*): evidence for complex infection history and disequilibrium distribution. Microb Ecol 65:731–739.

Zhao, D.-X., X.-F. Zhang, D.-S. Chen, Y.-K. Zhang, and X.-Y. Hong. 2013. *Wolbachia*-host interactions: host mating patterns affect *Wolbachia* density dynamics. PLoS One 8:e66373.

Zhou, P., X. Z. He, C. Chen, and Q. Wang. 2020. No evidence for inbreeding depression and inbreeding avoidance in a haplodiploid mite *Tetranychus ludeni* Zacher. Syst Appl Acarol 25:1723–1728.

