## Supplementary Materials for "*Wolbachia* strengthens the match between pre-mating and early post-mating isolation in spider mites"

### Box S1. Detailed information relating to the populations of *T. urticae* used in this study.

All information concerning the populations used in the experiments, gathered throughout our previous studies [1-4], is provided here and summarized in Table I.

DNA extractions from pools of 100 females for each population, and subsequent sequencing of a fragment of the nuclear ribosomal DNA *ITS2* (internal transcribed spacer 2) region and of the mitochondrial DNA Cytochrome Oxidase subunit I (*COI*), were performed as described in [4]. The results revealed that the two populations differ by 1 SNP in the ITS2 sequence and 21 SNPs in the COI sequence (genetic distance of 0.056 with Kimura 2-parameter; rate of variation gamma distribution; shape parameter = 1; [1]), indicating little genetic differentiation between the two populations despite almost complete postygotic isolation [1] (but see [5] for genomic differentiation between other populations of the two colour forms based on whole genome sequencing).

Although the prevalence of *Wolbachia* infection in the green population was not assessed upon field collection, it was of 100% in the red population [4], and, upon laboratory rearing, both populations were confirmed to be stably and fully infected with *Wolbachia*, but uninfected by any other known endosymbiont in spider mites (*Cardinium*, *Rickettsia*, *Spiroplasma* and *Arsenophonus*) [1, 3]. Multilocus sequence typing for *Wolbachia* (MLST; [6]), using primers and protocols described in [3], revealed that the red population is infected by the *Wolbachia* strain ST481, and the green population by the strain ST280 [2], both belonging to the *Ori* subgroup of the *Wolbachia* supergroup B [7]. These two strains are thus nearly identical, differing only in 1 SNP on the fbpA and coxA sequences of the MLST system [2], despite their striking difference in CI induction [1]. Both *Wolbachia*-infected populations, Ri and Gi, were subsequently treated with antibiotics as described below, to obtain their uninfected counterparts, Ru and Gu, respectively. Then, pools of 100 females from all populations were checked by multiplex PCR as described in [8] to confirm their *Wolbachia* infection status before performing the experiments.

The two populations used in the present study were obtained from treatments performed for the Experiment 1 of the previous Cruz et al. study [1]. These were done in November 2013 and February 2014 for the Gu and Ru populations, respectively. In both cases, a tetracycline solution (0.1%, w/v) was used for three successive generations as described in [3]. The same standard conditions as population rearing were used: same host plant (*Phaseolus vulgaris*, cv. Contender seedlings obtained from Germisem, Oliveira do Hospital, Portugal), temperature (24 ± 2°C), and photoperiod (16/8h L/D). The two *Wolbachia*-free populations were then maintained without antibiotics in the same mass-rearing conditions as the *Wolbachia*-infected populations for more than thirty generations, thereby avoiding potential side-effects of antibiotics [9-11]. This also ensured that spider mites at least partially recovered their microbiome beyond *Wolbachia* removal, despite that antibiotic treatments, or even the loss of *Wolbachia* itself, necessarily result in some permanent changes in the microbiome composition [12]. Yet, this recovery time has been proven effective in eliminating potential effects of the treatment on mite offspring production [3] or mating behaviour [13].

**Table I. Spider mite populations and *Wolbachia* infection.**

| **Population** | | **Ri** | **Gi** |
|---|---|---|---|
| **Host information** | **Original name** | AMP | TOM |
| | **Form** | red | green |
| | **Collection date** | 18/11/2013 | --/05/2010 |
| | **Host plant** | *Datura stramonium* | *Solanum lycopersicum* |
| | **Location** | Aldeia da Mata Pequena | Carregado |
| | **Coordinates** | 38.534363, -9.191163 | 39.078962, -8.993656 |
| | **Initial number** | 65♀ | 300♀ |
| | **ITS2**[1] | GU565314 | AM408031 |
| | **COI**[1] | MF428440 | HM486513 |
| ***Wolbachia* information** | **Isolate (id)**[2] | Turt_B_wUrtAmp (1858) | Turt_B_wUrtTom (1857) |
| | **Strain**[2] | 491 | 280 |
| | ***gatB* allele**[2] | 9 | 9 |
| | ***coxA* allele**[2] | 38 | 164 |
| | ***hcpA* allele**[2] | 143 | 143 |
| | ***ftsZ* allele**[2] | 23 | 23 |
| | ***fbpA* allele**[2] | 444 | 4 |
| | ***wsp*[1]** | GU014541 | GU014541 |
| | **CI level** | 57±3% | 1.3±0.5% |
| **References** | | [3, 4] | [1, 2, 14] |

[1] GenBank accession number matching with 100% coverage and identity at the nucleotide level.

[2] id, strain or allele number in the PubMLST *Wolbachia* MLST database, available at http://www.pubmlst.org/wolbachia/

**References**

1. Cruz, M. A., S. Magalhães, É. Sucena, and F. Zélé. 2021. *Wolbachia* and host intrinsic reproductive barriers contribute additively to postmating isolation in spider mites. Evolution 75:2085–2101.

2. Zélé, F., M. Altıntaş, I. Santos, I. Cakmak, and S. Magalhães. 2020. Population-specific effect of Wolbachia on the cost of fungal infection in spider mites. Ecol Evol 10:3868–80.

3. Zélé, F., I. Santos, M. Matos, M. Weill, F. Vavre, and S. Magalhães. 2020. Endosymbiont diversity in natural populations of Tetranychus mites is rapidly lost under laboratory conditions. Heredity 124:603–617.

4. Zélé, F., I. Santos, I. Olivieri, M. Weill, O. Duron, and S. Magalhães. 2018. Endosymbiont diversity and prevalence in herbivorous spider mite populations in South-Western Europe. FEMS Microbiol Ecol 94:fiy015.

5. Xue, W. X., J. T. Sun, J. Witters, M. Vandenhole, M. Dermauw, S. A. Bajda, E. A. Simma, N. Wybouw, E. Villacis-Perez, and T. Van Leeuwen. 2023. Incomplete reproductive barriers and genomic differentiation impact the spread of resistance mutations between green-and red-colour morphs of a cosmopolitan mite pest. Mol Ecol 32:4278-4297.

6. Baldo, L., J. C. D. Hotopp, K. A. Jolley, S. R. Bordenstein, S. A. Biber, R. R. Choudhury, C. Hayashi, M. C. J. Maiden, H. Tettelin, and J. H. Werren. 2006. Multilocus sequence typing system for the endosymbiont Wolbachia pipientis. Appl Environ Microbiol 72:7098–7110.

7. Zhang, Y.-K., X.-L. Ding, K.-J. Zhang, and X.-Y. Hong. 2013. *Wolbachia* play an important role in affecting mtDNA variation of *Tetranychus truncatus* (Trombidiformes: Tetranychidae). Environ Entomol 42:1240–1245.

8. Zélé, F., M. Weill, and S. Magalhães. 2018. Identification of spider-mite species and their endosymbionts using multiplex PCR. Exp Appl Acarol 74:123–138.

9. Ballard, J. W. O., and R. G. Melvin. 2007. Tetracycline treatment influences mitochondrial metabolism and mtDNA density two generations after treatment in *Drosophila*. Insect Mol Biol 16:799–802.

10. Zeh, J. A., M. M. Bonilla, A. J. Adrian, S. Mesfin, and D. W. Zeh. 2012. From father to son: transgenerational effect of tetracycline on sperm viability. Sci Rep 2:375.

11. O'Shea, K. L., and N. D. Singh. 2015. Tetracycline-exposed *Drosophila melanogaster* males produce fewer offspring but a relative excess of sons. Ecol and Evol 5:3130–3139.

12. Zhu, Y.-X., Y.-L. Song, A. A. Hoffmann, P.-Y. Jin, S.-M. Huo, and X.-Y. Hong. 2019. A change in the bacterial community of spider mites decreases fecundity on multiple host plants. Microbiologyopen 8:e00743.

13. Rodrigues, L. R., F. Zélé, I. Santos, and S. Magalhães. 2022. No evidence for the evolution of mating behavior in spider mites due to *Wolbachia*-induced cytoplasmic incompatibility. Evolution 76:623–635.

14. Clemente, S. H., L. R. Rodrigues, R. Ponce, S. A. M. Varela, and S. Magalhães. 2016. Incomplete species recognition entails few costs in spider mites, despite first-male precedence. Behav Ecol Sociobiol 70:1161–1170.

**Figure S1. Raw data for offspring production in the no-choice test.** Boxplot and individual data points for (a) daily oviposition (*i.e.*, number of eggs laid/number of days ovipositing), (b) number of unhatched eggs, (c) number of dead juveniles, (d) number of adult daughters, and (e) number of adult sons, produced by females from all types of crosses. Intra- and inter-population crosses were first performed between *Wolbachia*-uninfected females ("♀" on the bottom level of the x-axis) and *Wolbachia*-infected or uninfected males ("1st ♂" on the middle level of the x-axis). Then, all females that had mated were given the chance to remate with an uninfected male from their own population ("2nd ♂" on the top level of the x-axis; "-" indicates that the females did not mate with the second male). Colour legend indicates the categories of crosses.

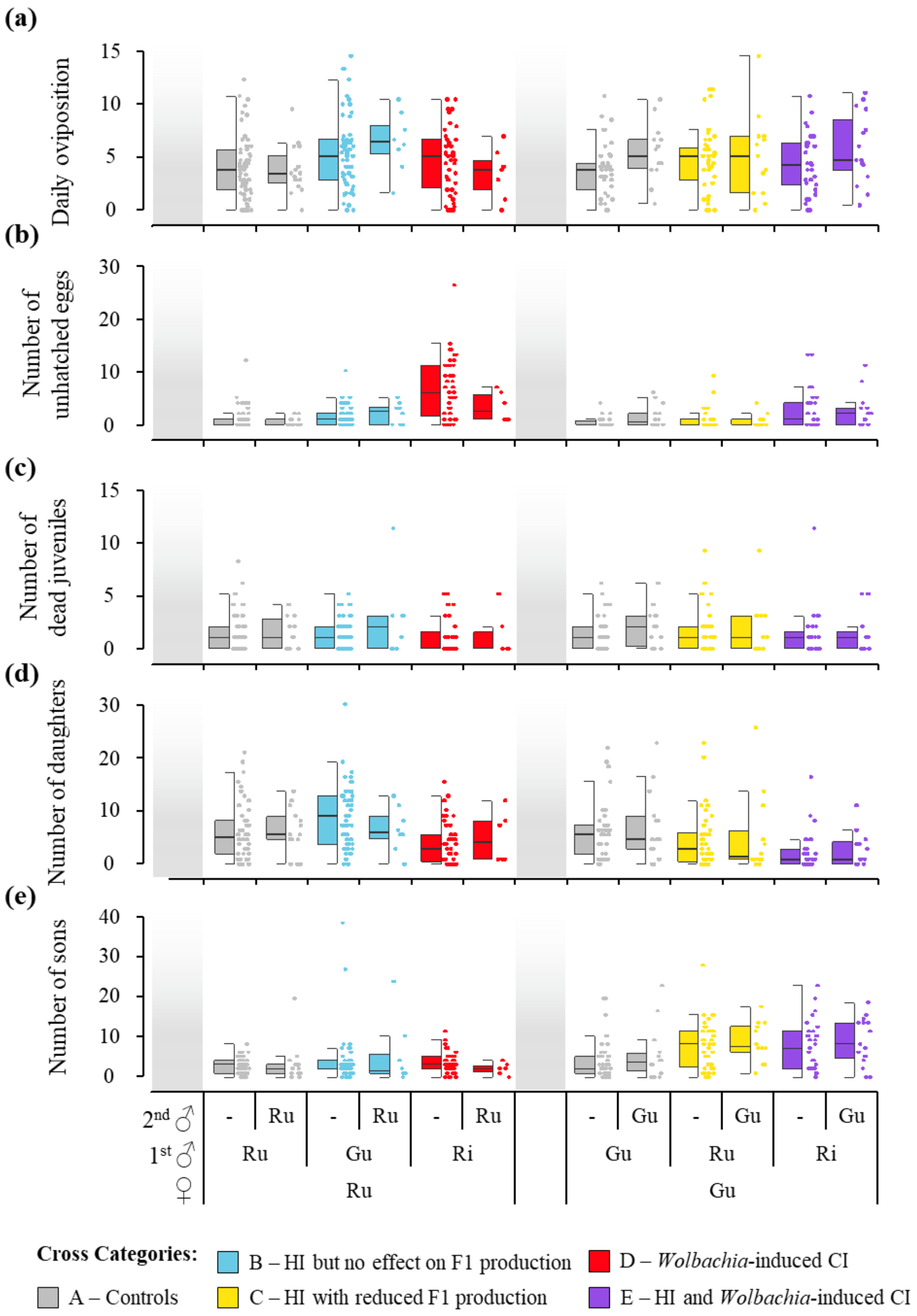

**Figure S2. Summary of the reproductive barriers occurring at different stages in life history of spider mites and variables measured in experimental tests to estimate the strength of each barrier ($RI_n$).** 'ns' means that the parameter was not used as the corresponding barrier was not found. Measured variables that may inform on a given barrier but does not allow unambiguous assessment are displayed between brackets. Those used to estimate each $RI_n$ parameter are displayed in bold.

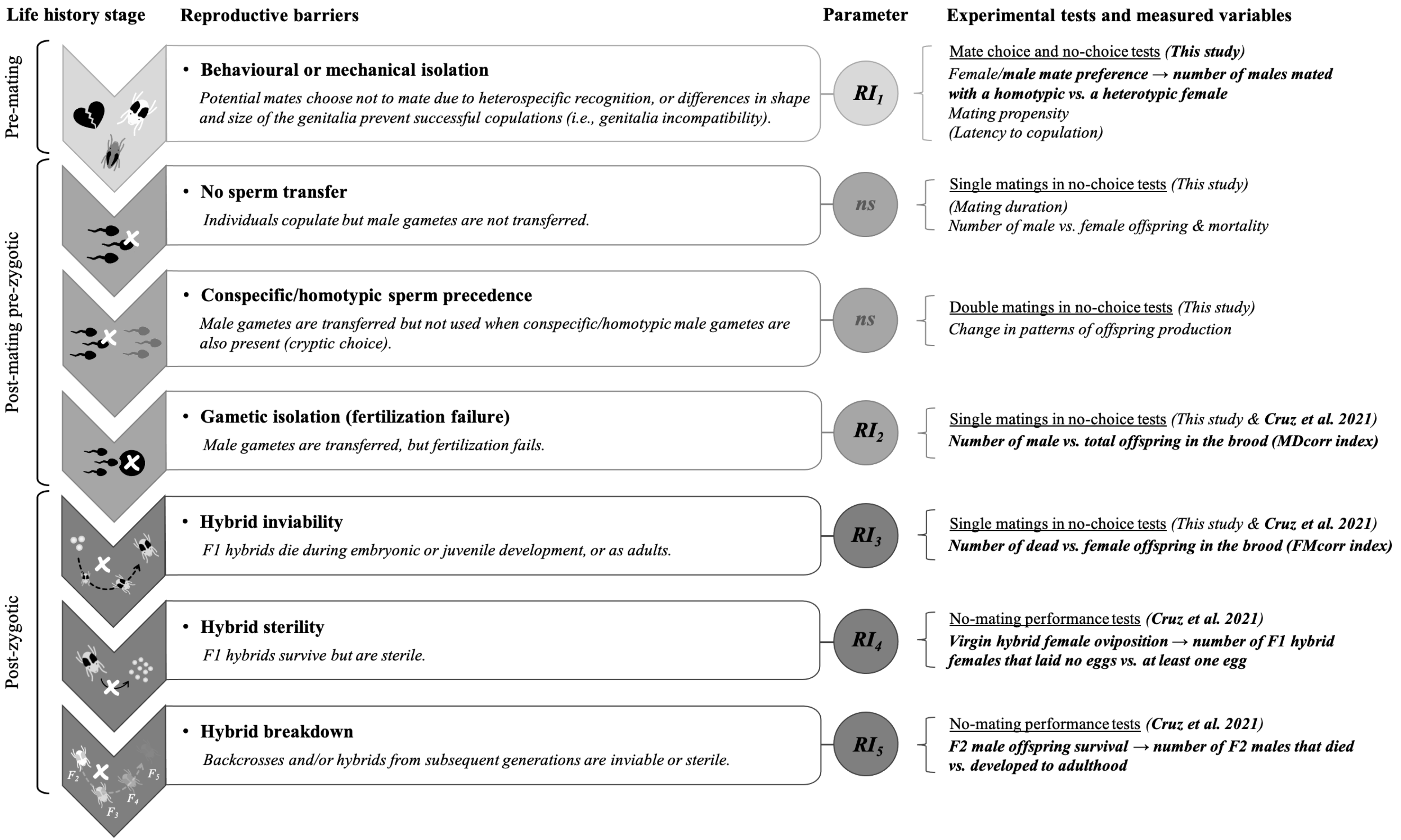

### Box S2. Complete description of the procedure used for all statistical analyses.

Latency to copulation and copulation duration were analysed using Cox proportional hazards mixed models (*coxme* procedure; coxme package), a non-parametric method traditionally used for survival analyses, but applicable to any time-to-event variable. This method, that analyses the likelihood of an event (beginning or end of a copulation) occurring over time, does not assume any particular error distribution [1]. All other data were analysed using generalized linear mixed models (*glmmTMB* procedure; glmmTMB package; [2]). Proportion data were computed either as binary response variables (*e.g.*, mated or not, chosen mate) or using a response variable generated by concatenating two dataset columns, binding together the number of successes and failures (*i.e.*, number of daughters *vs*. number of eggs – number of daughters in the analyses of female proportion) with the function *cbind*, to account for the total number of observations (*e.g.* the total number of eggs in the analyses of female proportion), thereby correcting for the fact that proportions calculated from low numbers of individuals are more prone to error [1]. For the same reason, we weighted each individual datum by the number of observations (*i.e.*, a "weights" argument was added to the models), in the analyses of the corrected variables $MD_{corr}$ and $FM_{corr}$, which are continuous variables bounded between 0 and 1. These data were subsequently analysed with a binomial or (zero-inflated) betabinomial error distribution when errors were overdispersed. Count data were analysed with a Poisson error distribution, and continuous data with a Gaussian, except for daily oviposition which was Box-Cox transformed ($\lambda$=0.549) to improve normality [1] and analysed with a zero-inflated Gaussian.

For the analyses of each response variable of the choice tests, the 'type' of the focal individual (*i.e.*, combination of population and infection status), and either the combination of provided mates (for the analyses of mating propensity and mate preference) or the chosen mate (for the analyses of latency to copulation and copulation duration), were fit as fixed explanatory variables, whereas the day and session (nested within day) of observation, the colour with which chosen males were painted (in the female choice test), and the combination of provided mates (for the analyses of latency to copulation and copulation duration) were fit as random explanatory variables. This model construction allowed testing for significant effects of the focal individual and of the available or chosen mates independently. In addition, to ascertain whether mate preferences differed from random choice in the analyses of mate preference, the intercept of the models was forced to zero to obtain the estimate of the fixed factor as the difference to a 0.5 probability (particularity of models with categorical factors and binomial distribution; [1]).

For the analyses of the no-choice test, mating propensity data obtained in the two mating events were analysed altogether, and the mating event was fit as fixed explanatory variable, along with the type of cross (*i.e.*, female x first male) and the interaction between these two factors, whereas the day and the session (nested within day) of observation were fit as random explanatory variables. This approach was used to determine whether females became less willing to mate with a second male following the first mating event. To avoid pseudoreplication in the analyses of copulation frequency, latency to copulation and copulation duration, data obtained in each mating event were analysed separately and only the type of cross was fit as fixed explanatory variable. Subsequently, to determine whether these response variables differed when females mated with the first or with the second male, the differences in copulation frequency, latency, and duration between 1$^{st}$ and 2$^{nd}$ mating events were computed for each individual female that mated at least once in both events. For that, the type of cross was fit as fixed explanatory variable, the day and session (nested within day) were fit as random explanatory variables, and the intercept was forced to zero to obtain the estimate of the fixed factor relative to no difference between mating events (*i.e.*, relative to zero). Finally, for the analyses of offspring data, the type of cross, the mating status of the females (*i.e.*, mated with only the first or both males), and the interaction between these two factors, were fit as fixed explanatory variables, whereas the day and session (nested within day) during which they mated were fit as random explanatory variables. This aimed at determining whether a second male was able to sire offspring following an incompatible cross (thereby reducing the negative effect of the latter), by comparing the offspring produced by females mated with one or with two males.

For all analyses, maximal models containing the complete set of explanatory variables were subsequently simplified by sequentially eliminating non-significant terms to establish a minimal model [1]. The significance of the explanatory variables was established using chi-square tests with the Anova function (car package; [3]). The significant values given in the text are for the minimal model, whereas non-significant values correspond to those obtained before deletion of the variable from the model [1]. When explanatory variables with more than two levels were found significant in the analyses of behavioural data, *a posteriori* contrasts between factor levels were carried out by aggregating factor levels together and testing the fit of the simplified model using a likelihood ratio test (function *anova;* [1]). When interactions between two explanatory variables were significant, the two variables were concatenated (*e.g.*, crosses and mating events) and *a posteriori* contrasts between the factor levels of the concatenated variable were carried out as described above. In all cases, Holm-Bonferroni corrections (*i.e.*, classical chi-squared Wald test for testing the global hypothesis $H_0$; [4]) were subsequently applied to account for multiple testing. Note that contrast analyses were not carried out for offspring production in the no-choice test because the results did not differ qualitatively from those of the previous study [5]. Finally, to determine the difference to random mating in the analyses of mate choice, as well as changes in mating behaviour between 1st and 2nd mating events in the no-choice test, coefficients (estimated as the difference with the zero-intercept) obtained from the maximal models for each combination of focal individual and pair of provided mates (mate choice test), or for each cross (no-choice test), were analysed with Z-tests using the function *test* (emmeans package; [6]).

**References**

1. Crawley, M. J. 2007. The R Book. John Wiley & Sons, Ltd, Chichester.
2. Brooks, M. E., K. Kristensen, K. J. van Benthem, A. Magnusson, C. W. Berg, A. Nielsen, H. J. Skaug, M. Mächler, and B. M. Bolker. 2017. glmmTMB balances speed and flexibility among packages for zero-inflated generalized linear mixed modeling. R J 9:378–400.
3. Fox, J., and S. Weisberg. 2019. An R Companion to Applied Regression. Third. Sage, Thousand Oaks.
4. Holm, S. 1979. A simple sequentially rejective multiple test procedure. Scand J Statist 6:65–70.
5. Cruz, M. A., S. Magalhães, É. Sucena, and F. Zélé. 2021. *Wolbachia* and host intrinsic reproductive barriers contribute additively to postmating isolation in spider mites. Evolution 75:2085–2101.
6. Lenth, R., H. Singmann, J. Love, P. Buerkner, and M. Herve. 2018. Emmeans: estimated marginal means, aka least-squares means.

**Table S1. Description of the statistical models used in analyses of the choice test.** The "data subset" column indicates the focal individuals included in each analysis, and the "sample size" column gives the corresponding number of observations. The "R subroutine" column indicates the type of model used for each analysis (b: binomial error structure; *coxme* is a non-parametric method that does not assume an error structure). Mating propensity and mate choice were analysed as binary response variables (*i.e.*, mated or not, and one mate chosen or the other, respectively). Latency to copulation and copulation duration were analysed as time to event variables using a Cox proportional hazards model (coxme), which did not include censored individuals as all non-mated focal individuals were excluded from these analyses. "Maximal model" gives the complete set of explanatory variables initially included in the model, from which non-significant fixed factors were sequentially removed to establish a minimal model. Round brackets indicate that the intercept of the variables was fitted as a random factor. **focal**: focal individual; **mates**: the combination of two mates presented to the focal individual; **chosen**: population and infection status of the chosen mate; **day/session**: the session, nested within the day, at which each observation was conducted; **colour**: the colour with which the chosen mate was painted.

| | Variable of interest | Response variable | Model No. | Data subset | Sample size | R subroutine | Maximal model | Minimal model |
|---|---|---|---|---|---|---|---|---|
| **Female behaviour** | Mating propensity | Females mated or not | 1.1 | All ♀ | 410 | glmmTMB [b] | focal + mates + (1\|day/session) | focal + (1\|day/session) |
| | Mate choice | Chosen male | 1.2 | Mated ♀ | 255 | glmmTMB [b] | 0 + focal + mates + (1\|colour) + (1\|day/session) | 0 + (1\|colour) + (1\|day/session) |
| | Latency to copulation | No. seconds before starting a copulation | 1.3 | Mated ♀ | 255 | coxme | focal + chosen + (1\|colour) + (1\|mates) + (1\|day/session) | 1+ (1\|colour) + (1\|mates) + (1\|day/session) |
| | Copulation duration | No. seconds spent copulating | 1.4 | Mated ♀ | 255 | coxme | focal + chosen + (1\|colour) + (1\|mates) + (1\|day/session) | focal + chosen + (1\|colour) + (1\|mates) + (1\|day/session) |
| **Male behaviour** | Mating propensity | Males mated or not | 1.5 | All ♂ | 425 | glmmTMB [b] | focal + mates + (1\|day/session) | mates + (1\|day/session) |
| | Mate choice | Chosen female | 1.6 | Mated ♂ | 274 | glmmTMB [b] | 0 + focal + mates + (1\|day/session) | 0 + mates + (1\|day/session) |
| | Latency to copulation | No. seconds before starting a copulation | 1.7 | Mated ♂ | 274 | coxme | focal + chosen + (1\|mates) + (1\|day/session) | 1 + (1\|mates) + (1\|day/session) |
| | Copulation duration | No. seconds spent copulating | 1.8 | Mated ♂ | 273[1] | coxme | focal + chosen + (1\|mates) + (1\|day/session) | focal + chosen + (1\|mates) + (1\|day/session) |

[1] Excludes one outlier that displayed an unusually high copulation duration of 19 minutes

**Table S2. Description of the statistical models used in analyses of the no-choice test.** The "data subset" column indicates the focal individuals that were included in each analysis, and the "sample size" column gives the corresponding number of individual crosses. The "R subroutine" column indicates the type of model used, with square brackets indicating the error structure used (b: binomial, p: Poisson, g: Gaussian, zig: zero-inflated Gaussian, bb: betabinomial, zibb: zero-inflated betabinomial; *coxme* is a non-parametric method that does not assume an error structure). Daily oviposition was Box-Cox transformed with $\lambda=0.549$ to improve the model fit. Cox proportional hazards models (coxme) used to analyse "time to event" variables (*i.e.* latency to copulation and copulation duration) do not include censored individuals as all non-mated focal individuals were excluded from these analyses. Models with a (beta)binomial error structure require either a binary response variable (*i.e.* mated or not for mating propensity), a concatenated response variable binding the number of successes and failures for a given outcome (*i.e.* being a female or not for female proportion), or a proportion bounded between 0 and 1 (for corrected indexes). In the latter case, a "weights" argument was added to the model to account for the number of observations per replicate. "Maximal model" gives the complete set of explanatory variables initially included in the model, from which non-significant fixed factors were sequentially removed to establish a minimal model. Round brackets indicate that the intercept of the variables was fitted as a random factor. **event**: the mating event (first or second) in which data was obtained; **cross**: combination between the populations of the female and the male in the first mating event; **day/session**: the session, nested within the day, at which each observation was conducted; **status**: whether the female mated with the male in the second mating event or not; **eggs**: total number of eggs laid; **days**: total number of days the female was alive; **$MD_{obs}$**: proportion of adult sons relative to total offspring; **CCMD**: mean proportion of adult sons relative to total offspring in control crosses; **$FM_{obs}$**: proportion of unhatched eggs relative to adult females; **CCFM**: mean proportion of unhatched eggs relative to females in control crosses; **daughters/sons**: total number of adult daughters/sons in each cross.

| | Variable of interest | Response variable | Model No. | Data subset | Sample size | R subroutine [family] | Maximal model | Minimal model |
|---|---|---|---|---|---|---|---|---|
| **Mating propensity and frequency** | Mating propensity | Females mated or not | 2.1 | Complete dataset | 639+369[1] | glmmTMB [b] | cross * event + (1\|day/session) | cross * event + (1\|day/session) |
| | Copulation frequency with the 1st ♂ | Number of copulations | 2.2 | ♀ mated with the 1st ♂ | 369 | glmmTMB [p] | cross + (1\|day/session) | 1 + (1\|day/session) |
| | Copulation frequency with the 2nd ♂ | Number of copulations | 2.3 | ♀ mated with the two ♂ | 72 | glmmTMB [p] | cross + (1\|day/session) | 1 + (1\|day/session) |
| | Difference in copulation frequency with the 1st and 2nd ♂ | Number of copulations with the 1st ♂ – number of copulations with the 2nd ♂ | 2.4 | ♀ mated with the two ♂ | 72 | glmmTMB [g] | 0 + cross + (1\|day/session) | 0 + (1\|day/session) |
| **Latency to copulation** | Latency to copulation with the 1st ♂ | Time to 1st copulation with the 1st ♂ | 2.5 | ♀ mated with the 1st ♂ | 369 | coxme | cross + (1\|day/session) | cross + (1\|day/session) |
| | Latency to copulation with the 2nd ♂ | Time to 1st copulation with the 2nd ♂ | 2.6 | ♀ mated with the two ♂ | 72 | coxme | cross + (1\|day/session) | 1 + (1\|day/session) |
| | Difference in latency to copulation with the 1st and 2nd ♂ | Time to 1st copulation with the 1st ♂ – Time to 1st copulation with the 2nd ♂ | 2.7 | ♀ mated with the two ♂ | 72 | glmmTMB [g] | 0 + cross + (1\|day/session) | 0 + (1\|day/session) |
| **Copulation duration** | Copulation duration with the 1st ♂ | ∑ duration of copulation(s) with the 1st ♂ | 2.8 | ♀ mated with the 1st ♂ | 369 | coxme | cross + (1\|day/session) | cross + (1\|day/session) |
| | Copulation duration with the 2nd ♂ | ∑ duration of copulation(s) with the 2nd ♂ | 2.9 | ♀ mated with the two ♂ | 71[2] | coxme | cross + (1\|day/session) | 1 + (1\|day/session) |
| | Difference in copulation duration with the 1st and 2nd ♂ | ∑ duration of copulation(s) with the 1st ♂ – ∑ duration of copulation(s) with the 2nd ♂ | 2.10 | ♀ mated with the two ♂ | 71[2] | glmmTMB [g] | 0 + cross + (1\|day/session) | 0 + cross + (1\|day/session) |
| **Offspring production** | Daily oviposition | $[(eggs/days)^{\lambda} – 1]/\lambda$ | 2.11 | All mated ♀ | 346[3] | glmmTMB [zig] | cross * status + (1\|day/session) | cross + (1\|day/session) |
| | Male overproduction ($MD_{corr}$) | $(MD_{obs}$-CCMD)/(1-CCMD) | 2.12 | All mated ♀ | 319[4] | glmmTMB [bb] | cross * status + (1\|day/session) | cross + (1\|day/session) |
| | Female mortality ($FM_{corr}$) | $(FM_{obs}$-CCFM)/(1-CCFM) | 2.13 | All mated ♀ | 296[5] | glmmTMB [bb] | cross * status + (1\|day/session) | cross + (1\|day/session) |
| | Female proportion (FP) | cbind(daughters,eggs-daughters) | 2.14 | All mated ♀ | 319[4] | glmmTMB [zibb] | cross * status + (1\|day/session) | cross + (1\|day/session) |

[1] Combines data obtained in the 1st and 2nd mating events (639 and 369 observed females, respectively)
[2] Excludes one outlier that displayed an unusually high copulation duration of over 14 minutes
[3] Includes all mated females (*i.e.*, mated only with the 1st male or mated with the two males)
[4] Includes all mated females that produced at least 1 egg
[5] Includes all mated females that produced at least 1 unhatched egg or 1 adult daughter

**Table S3. Mating propensity and mate choice observed in the choice test.** The mean (± s.e.) proportion of focal males and females that mated with one of the proposed mates ('Mating propensity' column), and of *Wolbachia*-infected or uninfected green or red form chosen mates ('Chosen proportion' column), are provided for each choice category (*cf.* description in Table 1). $N_{mated}$: Number of focal individuals that mated with either of the proposed mates. $N_{total}$: total number of tested focal individuals, including those that did not mate. Ru: red uninfected; Ri: red infected; Gu: green uninfected; Gi: green infected.

| | Category | Focal individual | Proposed mates | Mating propensity (%) | Chosen mate | Chosen proportion (%) | $N_{mated}$ | $N_{total}$ |
|---|---|---|---|---|---|---|---|---|
| Female choice test | I | Ru | Ru *vs* Ri | 73.08 ± 6.21 | Ru | 45.95 ± 8.31 | 17 | 52 |
| | | | | | Ri | 54.05 ± 8.31 | 20 | |
| | | Gu | Gu *vs* Gi | 62.75 ± 6.84 | Gu | 61.29 ± 8.89 | 19 | 51 |
| | | | | | Gi | 38.71 ± 8.89 | 12 | |
| | II | Ru | Ru *vs* Gu | 61.54 ± 6.81 | Ru | 48.39 ± 9.12 | 15 | 52 |
| | | | | | Gu | 51.61 ± 9.12 | 16 | |
| | | Gu | Ru *vs* Gu | 56.86 ± 7.00 | Ru | 34.48 ± 8.98 | 10 | 51 |
| | | | | | Gu | 65.52 ± 8.98 | 19 | |
| | III | Ri | Ri *vs* Gi | 72.55 ± 6.31 | Ri | 55.56 ± 8.40 | 20 | 51 |
| | | | | | Gi | 44.44 ± 8.40 | 16 | |
| | | Gi | Ri *vs* Gi | 50.98 ± 7.07 | Ri | 46.15 ± 9.97 | 12 | 51 |
| | | | | | Gi | 53.85 ± 9.97 | 14 | |
| | IV | Ru | Ri *vs* Gi | 68.63 ± 6.56 | Ri | 31.43 ± 7.96 | 11 | 51 |
| | | | | | Gi | 68.57 ± 7.96 | 24 | |
| | | Gu | Ri *vs* Gi | 50.98 ± 7.07 | Ri | 60.00 ± 10.00 | 15 | 51 |
| | | | | | Gi | 40.00 ± 10.00 | 10 | |
| Male choice test | I | Ri | Ru *vs* Ri | 69.81 ± 3.67 | Ru | 40.54 ± 8.18 | 15 | 53 |
| | | | | | Ri | 59.46 ± 8.18 | 22 | |
| | | Gi | Gu *vs* Gi | 39.62 ± 6.78 | Gu | 47.62 ± 11.17 | 10 | 53 |
| | | | | | Gi | 52.38 ± 11.17 | 11 | |
| | II | Ru | Ru *vs* Gu | 61.11 ± 6.70 | Ru | 63.63 ± 8.50 | 21 | 54 |
| | | | | | Gu | 36.36 ± 8.50 | 12 | |
| | | Gu | Ru *vs* Gu | 64.15 ± 6.65 | Ru | 79.41 ± 7.04 | 27 | 53 |
| | | | | | Gu | 20.59 ± 7.04 | 7 | |
| | III | Ri | Ri *vs* Gi | 73.58 ± 6.11 | Ri | 74.36 ± 7.08 | 29 | 53 |
| | | | | | Gi | 25.64 ± 7.08 | 10 | |
| | | Gi | Ri *vs* Gi | 64.15 ± 6.65 | Ri | 79.41 ± 7.04 | 27 | 53 |
| | | | | | Gi | 20.59 ± 7.04 | 7 | |
| | IV | Ri | Ru *vs* Gu | 77.36 ± 5.80 | Ru | 80.49 ± 6.27 | 33 | 53 |
| | | | | | Gu | 19.51 ± 6.27 | 8 | |
| | | Gi | Ru *vs* Gu | 66.04 ± 6.57 | Ru | 80.00 ± 6.86 | 28 | 53 |
| | | | | | Gu | 20.00 ± 6.86 | 7 | |

**Table S4. Results of the contrast analyses performed among or between populations of focal individuals/combination of mates that significantly affected a given behavioural variable in the female (a) and male (b) choice tests.** Holm corrections were used to account for multiple testing. Ru: red uninfected; Ri: red infected; Gu: green uninfected; Gi: green infected.

| **(a) Contrasts for the female choice test** | **Chi-value** | **Df** | **P-value** |
|---|---|---|---|
| **Among female populations (regardless of provided males) for mating propensity (Fig. 1a)** | | | |
| Among red ♀, regardless of *Wolbachia* infection (within group "a") | 0.31 | 1 | 0.78 |
| Among green ♀, regardless of *Wolbachia* infection (within group "b") | 0.75 | 1 | 0.78 |
| Red ♀ *vs*. green ♀ (group "a" *vs* "b") | 9.07 | 1 | 0.008 |
| **Among male populations for copulation duration (Fig. 2c)** | | | |
| Among green ♂ (Gi *vs* Gu; within group "b") | 0.09 | 1 | 0.76 |
| Ri ♂ *vs* Ru ♂ (group "a" *vs* "ab") | 0.99 | 1 | 0.64 |
| Ru ♂ *vs* green ♂ (group "ab" *vs* "b") | 2.40 | 1 | 0.36 |
| Ri ♂ *vs* green ♂ (group "a" *vs* "b") | 6.10 | 1 | 0.05 |
| **Among female populations for copulation duration (Fig. 2d)** | | | |
| Between uninfected ♀ (Ru *vs* Gu; within group "a") | 1.08 | 1 | 0.64 |
| Uninfected ♀ *vs* Gi ♀ (group "a" *vs* "ab") | 1.50 | 1 | 0.64 |
| Gi ♀ *vs* Ri ♀ (group "ab" *vs* "b") | 1.56 | 1 | 0.64 |
| Uninfected ♀ *vs* Ri ♀ (group "a" *vs* "b") | 7.66 | 1 | 0.02 |

| **(b) Contrasts for the male choice test** | **Chi-value** | **Df** | **P-value** |
|---|---|---|---|
| **Among males provided with different females for mating propensity (Fig. 1c)** | | | |
| Among all ♂ except Gi ♂ provided with Gi and Gu ♀ (within group "a") | 0.12 | 2 | 0.99 |
| Gi ♂ provided with Gi and Gu ♀ *vs*. all other ♂ (group "a" *vs* "b") | 15.61 | 1 | <0.001 |
| **Among males provided with different females for mate choice (Fig. 1d)** | | | |
| Among ♂ provided with 'homotypic *vs* heterotypic' ♀ (within group "a") | 0.28 | 1 | 1 |
| Among ♂ provided with 'infected *vs* uninfected' ♀ (within group "b") | 0.37 | 1 | 1 |
| Between ♂ provided with 'homotypic *vs* heterotypic' and 'infected *vs* uninfected' ♀ (group "a" *vs* "b") | 9.35 | 1 | 0.007 |
| **Among male populations for copulation duration (Fig. 2c)** | | | |
| Among red ♂ (Ri *vs* Ru; within group "a") | 1.44 | 1 | 0.46 |
| Among green ♂ (Gi *vs* Gu; within group "b") | 0.52 | 1 | 0.47 |
| Red ♂ *vs* green ♂ (group "a" *vs* group "b") | 24.87 | 1 | <0.0001 |
| **Among female populations for copulation duration (Fig. 2d)** | | | |
| Among uninfected ♀ (Ru *vs* Gu; within group "a") | 3.22 | 1 | 0.15 |
| Among infected ♀ (Ri *vs* Gi; within group "b") | 0.01 | 1 | 0.92 |
| Uninfected ♀ *vs* infected ♀ (group "a" *vs* "b") | 21.67 | 1 | <0.0001 |

**Table S5. Effect size for *Wolbachia* infection and/or colour form of the provided mates on the choice of focal individuals in the choice test.** Regression coefficients (β), standard errors (SE), t ratios and P-values associated to the difference with a 50/50 choice were obtained from t-tests performed on models 1.2 and 1.6 (*cf.* Table S1), in which the intercept was set to zero. Grey cells highlight significant differences at the 5% level. Ru: red uninfected; Ri: red infected; Gu: green uninfected; Gi: green infected.

| | Category | Focal individual | Proposed mates | β | SE (β) | t ratio | P-value |
|---|---|---|---|---|---|---|---|
| Female choice test | I | Ru | Ru *vs* Ri | 0.148 | 0.340 | 0.434 | 0.665 |
| | | Gu | Gu *vs* Gi | -0.458 | 0.377 | -1.212 | 0.227 |
| | II | Ru | Ru *vs* Gu | -0.067 | 0.370 | -0.182 | 0.856 |
| | | Gu | Ru *vs* Gu | -0.664 | 0.401 | -1.656 | 0.099 |
| | III | Ri | Ri *vs* Gi | 0.227 | 0.345 | 0.658 | 0.511 |
| | | Gi | Ri *vs* Gi | -0.146 | 0.403 | -0.363 | 0.717 |
| | IV | Ru | Ri *vs* Gi | -0.789 | 0.373 | -2.114 | 0.036 |
| | | Gu | Ri *vs* Gi | 0.421 | 0.420 | 1.003 | 0.317 |
| Male choice test | I | Ri | Ru *vs* Ri | 0.381 | 0.338 | 1.128 | 0.260 |
| | | Gi | Gu *vs* Gi | 0.104 | 0.443 | 0.234 | 0.815 |
| | II | Ru | Ru *vs* Gu | 0.564 | 0.366 | 1.541 | 0.125 |
| | | Gu | Ru *vs* Gu | 1.350 | 0.427 | 3.162 | 0.002 |
| | III | Ri | Ri *vs* Gi | 1.069 | 0.370 | 2.886 | 0.004 |
| | | Gi | Ri *vs* Gi | 1.356 | 0.428 | 3.168 | 0.002 |
| | IV | Ri | Ru *vs* Gu | 1.419 | 0.397 | 3.576 | 0.0004 |
| | | Gi | Ru *vs* Gu | 1.399 | 0.431 | 3.246 | 0.001 |

**Table S6. Latency to copulation and copulation duration observed in the choice test.** Mean (± s.e.) latency to copulation and copulation duration (in seconds) of matings that occurred between *Wolbachia*-infected or uninfected green or red males and females, independently of the combination of proposed mates. $N_{mated}$: Number of focal individuals that mated with each type of mate, regardless of the combination of proposed mates. Ru: red uninfected; Ri: red infected; Gu: green uninfected; Gi: green infected.

| | Focal individual | Chosen mate | Latency to copulation (seconds) | Copulation duration (seconds) | $N_{mated}$ |
|---|---|---|---|---|---|
| Female choice test | Ru | Ru | 494.87 ± 64.23 | 221.29 ± 16.90 | 32 |
| | | Ri | 463.29 ± 67.80 | 222.94 ± 12.32 | 32 |
| | | Gu | 372.17 ± 91.97 | 262.56 ± 15.94 | 17 |
| | | Gi | 506.73 ± 66.98 | 245.42 ± 17.83 | 24 |
| | Ri | Ri | 636.18 ± 87.92 | 195.85 ± 14.23 | 21 |
| | | Gi | 429.75 ± 77.92 | 221.41 ± 16.45 | 16 |
| | Gu | Gu | 631.25 ± 74.31 | 263.00 ± 12.84 | 38 |
| | | Gi | 526.85 ± 86.13 | 270.82 ± 15.77 | 23 |
| | | Ru | 650.75 ± 138.97 | 229.44 ± 22.08 | 10 |
| | | Ri | 545.38 ± 94.27 | 233.24 ± 22.07 | 16 |
| | Gi | Gi | 475.52 ± 120.38 | 251.51 ± 12.43 | 14 |
| | | Ri | 469.98 ± 74.93 | 195.42 ± 25.61 | 12 |
| Male choice test | Ru | Ru | 497.76 ± 97.10 | 192.92 ± 18.96 | 21 |
| | | Gu | 460.00 ± 65.06 | 219.81 ± 11.86 | 12 |
| | Ri | Ru | 428.09 ± 61.59 | 224.38 ± 10.87 | 48 |
| | | Ri | 366.29 ± 48.21 | 191.50 ± 7.75 | 51 |
| | | Gu | 536.82 ± 151.91 | 206.52 ± 22.98 | 8 |
| | | Gi | 323.74 ± 74.54 | 175.01 ± 18.39 | 10 |
| | Gu | Ru | 379.36 ± 73.41 | 264.95 ± 17.98 | 27 |
| | | Gu | 665.48 ± 165.43 | 261.72 ± 27.94 | 7 |
| | Gi | Ru | 445.83 ± 61.79 | 269.21 ± 16.32 | 28 |
| | | Ri | 449.31 ± 72.70 | 216.92 ± 11.13 | 27 |
| | | Gu | 330.29 ± 43.30 | 229.36 ± 15.68 | 17 |
| | | Gi | 547.45 ± 80.19 | 219.25 ± 16.89 | 18 |

**Table S7. Mating behaviour and offspring production in each cross performed in the no-choice test.** Intra- and inter-population crosses were first performed ('1st mating event') between uninfected red or green females (Ru or Gu), and *Wolbachia*-infected red (Ri), uninfected red (Ru), or uninfected green (Gu) males. Then, mated females were presented with an uninfected male from their own population ('2nd mating event'). $N_{total}$, $N_{mated}$ and $N_{remated}$ correspond, respectively, to the total number of tested pairs, mated pairs during the first mating event, and mated pairs during the second mating event. Copulation frequency corresponds to the mean (± s.e.) number of copulations occurring with the same male. Latency to copulation corresponds to the mean (± s.e.) time in seconds before a first copulation started. Copulation duration corresponds to the mean (± s.e.) cumulative time in seconds the individuals spent copulating. Daily oviposition corresponds to the mean (± s.e.) number of eggs laid by a female, divided by the number of days the female was alive. Male overproduction and female mortality correspond, respectively, to the mean (± s.e.) increase in the proportion of adult male offspring and unhatched female eggs, relative to control crosses (*cf.* formula for the $MD_{corr}$ and $FM_{corr}$ indexes in the main text). Female proportion corresponds to the percentage of female offspring in the brood. For each tested behavioural variable (*i.e.*, each row, but those relating to the offspring), identical or absent superscripts indicate nonsignificant differences between crosses at the 5% level. As it was already the focus of our previous study (Cruz et al. 2021), contrast analyses to test for differences among crosses were not performed for $MD_{corr}$, $FM_{corr}$ and FP. Ru: red uninfected; Ri: red infected; Gu: green uninfected; Gi: green infected.

| **Female** | | **Ru** | | | **Gu** | | |
|---|---|---|---|---|---|---|---|
| | **1st Male** | **Ru** | **Ri** | **Gu** | **Gu** | **Ru** | **Ri** |
| **1st mating event** | **$N_{mated}$/$N_{total}$ (Mating propensity)** | 73/117 (62.39%)[a] | 58/101 (57.43%)[ab] | 66/100 (66.00%)[a] | 59/118 (50.00%)[b] | 65/103 (63.11%)[a] | 48/100 (48.00%)[b] |
| | **Copulation frequency[1]** | 2.10 ± 0.18 | 1.86 ± 0.18 | 2.39 ± 0.18 | 2.25 ± 0.24 | 1.94 ± 0.15 | 1.73 ± 0.19 |
| | **Latency to copulation[1]** | 917.74 ± 110.33[ab] | 810.34 ± 104.17[a] | 790.59 ± 79.48[a] | 1178.53 ± 119.23[b] | 1082.23 ± 107.86[b] | 1201.52 ± 158.42[b] |
| | **Copulation duration[1]** | 257.34 ± 30.08[a] | 244.86 ± 21.79[a] | 283.98 ± 8.58[b] | 270.46 ± 15.75[b] | 230.20 ± 13.21[a] | 214.98 ± 10.21[a] |
| | **Daily oviposition[2]** | 4.27 ± 0.42 | 5.03 ± 0.46 | 5.63 ± 0.49 | 3.81 ± 0.41 | 4.93 ± 0.48 | 4.59 ± 0.56 |
| | **Male overproduction ($MD_{corr}$)[2]** | 9.53 ± 3.30 | 9.38 ± 3.92 | 7.60 ± 2.74 | 15.17 ± 4.65 | 37.28 ± 5.50 | 34.00 ± 7.03 |
| | **Female mortality ($FM_{corr}$)[2]** | 17.35 ± 4.65 | 54.57 ± 5.56 | 2.94 ± 1.51 | 7.33 ± 3.47 | 21.74 ± 6.58 | 43.69 ± 8.85 |
| | **Female proportion (FP)[2]** | 49.28 ± 3.72 | 23.83 ± 3.38 | 60.86 ± 3.04 | 52.81 ± 4.59 | 31.88 ± 4.83 | 26.37 ± 5.81 |
| | **2nd Male** | | **Ru** | | | **Gu** | |
| **2nd mating event** | **$N_{remated}$/$N_{mated}$ (Mating propensity)** | 15/73 (20.55%) | 7/58 (12.07%) | 8/66 (12.12%) | 14/59 (23.73%) | 13/65 (20.00%) | 15/48 (31.25%) |
| | **Copulation frequency[3]** | 1.33 ± 0.16 | 1.71 ± 0.42 | 1.75 ± 0.41 | 1.71 ± 0.40 | 1.62 ± 0.27 | 1.40 ± 0.13 |
| | **Latency to copulation[3]** | 1821.80 ± 243.53 | 1084.71 ± 380.41 | 1634.00 ± 329.01 | 1966.07 ± 230.15 | 1292.39 ± 258.26 | 1437.67 ± 199.63 |
| | **Copulation duration[3]** | 92.27 ± 18.46 | 103.00 ± 31.11 | 94.43 ± 22.46 | 104.43 ± 12.92 | 134.54 ± 24.96 | 109.33 ± 17.90 |
| | **Daily oviposition[3]** | 4.02 ± 0.64 | 3.62 ± 0.96 | 6.77 ± 1.04 | 5.69 ± 0.75 | 5.37 ± 1.16 | 6.12 ± 0.91 |
| | **Male overproduction ($MD_{corr}$)[3]** | 21.43 ± 11.38 | 0.61 ± 0.61 | 11.41 ± 8.78 | 14.02 ± 6.01 | 42.46 ± 6.97 | 40.48 ± 8.18 |
| | **Female mortality ($FM_{corr}$)[3]** | 3.73 ± 2.17 | 41.65 ± 15.45 | 24.41 ± 12.01 | 19.88 ± 9.75 | 17.50 ± 11.15 | 45.47 ± 12.22 |
| | **Female proportion (FP)[3]** | 54.67 ± 8.76 | 38.42 ± 9.34 | 41.96 ± 10.03 | 45.13 ± 8.16 | 26.23 ± 6.84 | 27.53 ± 8.87 |

[1] Includes all females that mated with the 1st male
[2] Only includes females that mated with the 1st male but not with the 2nd male
[3] Only includes females that mated with both males

Cruz, M. A., S. Magalhães, É. Sucena, and F. Zélé. 2021. *Wolbachia* and host intrinsic reproductive barriers contribute additively to postmating isolation in spider mites. Evolution. 75:2085–210

**Table S8. Results of the contrast analyses performed among or between different types of crosses for the mating propensities observed in the no-choice test (Fig. 3).** Holm corrections were used to account for multiple testing. Ru: red uninfected; Ri: red infected; Gu: green uninfected; Gi: green infected.

| Contrasts performed | Chi-value | Df | P-value |
|---|---|---|---|
| **Between types of crosses within the 1st mating event** | | | |
| Among mating propensities observed in the 1st mating event for all types of crosses, but that of Gu ♀ presented with Ri or Gu ♂ (within group "a") | 1.70 | 3 | 0.77 |
| Among mating propensities observed in the 1st mating event for Ru and Gu ♀ presented with Ri ♂ and Gu ♀ presented with Gu ♂ (within group "b") | 1.91 | 2 | 0.77 |
| Between mating propensities observed in the 1st mating event for Ru ♀ presented with Ru or Gu ♂ or Gu ♀ presented with Ru ♂, and those observed for Gu ♀ presented with Gu or Ri ♂ (group "a" *vs* "bc") | 11.36 | 1 | 0.004 |
| **Between types of crosses within the 2nd mating event** | | | |
| Among all mating propensities observed in the 2nd mating event (within group "d") | 9.41 | 5 | 0.28 |
| **Between the two mating events within each type of cross** | | | |
| Between the mating propensities observed in the 1st event for Gu ♀ presented with Gu or Ri ♂, and the mating propensity observed in the 2nd event for Gu ♀ that had mated with Ri ♂ in the 1st event (within group "c") | 6.50 | 2 | 0.16 |
| Between the mating propensity observed in the 1st event for Gu ♀ presented with Gu or Ri ♂ and all mating propensities observed in the 2nd event but that of Gu ♀ that had mated with Ri ♂ in the 1st event (group "bc" *vs* "d") | 64.26 | 1 | <0.0001 |

**Table S9. Results of the contrast analyses performed among or between types of crosses for latency to copulation and copulation duration in the 1st mating event of the no-choice test (Fig. 4).** Holm corrections were used to account for multiple testing. Ru: red uninfected; Ri: red infected; Gu: green uninfected; Gi: green infected.

| Contrasts performed | Chi-value | Df | P-value |
|---|---|---|---|
| **Among types of crosses for latency to copulation (Fig. 4a)** | | | |
| Ru ♀ mated with Gu ♂ *vs* mated with Ri ♂ (within group "a") | 0.30 | 1 | 1 |
| Among Gu ♀, regardless of their mates (within group "b") | 0.92 | 2 | 1 |
| Ru ♀ mated with Gu or Ri ♂ *vs* mated with Ru ♂ (group "a" *vs* "ab") | 1.43 | 1 | 0.70 |
| Gu ♀ mated with any male *vs* Ru ♀ mated with Ru ♂ (group "b" *vs* "ab") | 2.60 | 1 | 0.43 |
| Ru ♀ mated with Gu or Ri ♂ *vs* Gu ♀ mated with any male (group "a" *vs* "b") | 11.51 | 1 | 0.003 |
| **Among male populations* for copulation duration (Fig. 4c)** | | | |
| Among red ♂ (within group "a") | 2.90 | 3 | 0.82 |
| Among green ♂ (within group "b") | 0.68 | 1 | 0.82 |
| Red ♂ *vs* green ♂ (group "a" *vs* group "b") | 17.42 | 1 | <0.0001 |

*regardless of female population or *Wolbachia* infection

**Table S10. Effect size for the difference in latency to copulation and copulation duration between the 1st and 2nd mating events in the no-choice test.** Mean differences (± SE), regression coefficients (β) and associated standard errors (SE) are provided for each type of cross. t ratios and P-values associated to the difference between the regression coefficients and the zero intercept (*cf.* Fig 4b and 4d) were obtained from t-tests performed on models 2.7 and 2.10 (*cf.* Table S2). Dark and light grey cells highlight significant effects at the 5% and 10% level, respectively. Ru: red uninfected; Ri: red infected; Gu: green uninfected; Gi: green infected.

| | Cross (♀ x 1st♂) | Mean ± SE | β | SE (β) | t ratio | P-value |
|---|---|---|---|---|---|---|
| Latency to copulation | Ru x Ru | 576.60 ± 360.41 | 577 | 349 | 1.655 | 0.103 |
| | Ru x Gu | 830.38 ± 404.62 | 833 | 478 | 1.743 | 0.086 |
| | Ru x Ri | 251.29 ± 630.96 | 244 | 511 | 0.478 | 0.635 |
| | Gu x Gu | 899.93 ± 307.37 | 899 | 361 | 2.490 | 0.015 |
| | Gu x Ru | -157.62 ± 408.85 | -158 | 375 | -0.422 | 0.674 |
| | Gu x Ri | 323.40 ± 408.24 | 324 | 349 | 0.928 | 0.357 |
| Copulation duration | Ru x Ru | -101.27 ± 31.14 | -101.1 | 31.6 | -3.201 | 0.002 |
| | Ru x Gu | -195.29 ± 45.64 | -197.1 | 44.7 | -4.412 | <0.0001 |
| | Ru x Ri | -51.86 ± 64.76 | -40.6 | 45.9 | -0.885 | 0.380 |
| | Gu x Gu | -137.57 ± 23.39 | -142.0 | 32.9 | -4.314 | 0.0001 |
| | Gu x Ru | -55.77 ± 43.09 | -48.3 | 33.9 | -1.424 | 0.159 |
| | Gu x Ri | -104.67 ± 27.94 | -100.9 | 32.2 | -3.136 | 0.003 |

**Table S11. Estimated strength of reproductive isolation (*$RI_n$*) and contribution (*$C_n$*) to total isolation (*$T$*) of each reproductive barrier identified within and between spider mite populations at each stage *$n$* of their life history.** Ru: red uninfected; Ri: red infected; Gu: green uninfected; Gi: green infected; NT: not tested; NA: not applicable.

| **Type of cross** | Strength /contribution (%) | **Reproductive barrier at stage *n*** | | | | | **Total isolation** |
|---|---|---|---|---|---|---|---|
| | | Assortative mating ($n$ = 1) | Fertilization failure ($n$ = 2) | Hybrid inviability ($n$ = 3) | Hybrid sterility ($n$ = 4) | Hybrid breakdown ($n$ = 5) | |
| Ru ♀ x Ri ♂ | $RI_n$ | 18.92 | 0 | 39.85 | 0 | 0 | 51.23 |
| | $C_n$ | 18.92 | 0 | 32.31 | 0 | 0 | |
| Gu ♀ x Gi ♂ | $RI_n$ | 4.76 | 0 | 0 | 1.03 | 0 | 5.74 |
| | $C_n$ | 4.76 | 0 | 0 | 0.98 | 0 | |
| Ru ♀ x Gu ♂ | $RI_n$ | 0* | 0 | 0 | 97.89 | 100 | 100 |
| | $C_n$ | 0 | 0 | 0 | 97.89 | 2.11 | |
| Gu ♀ x Ru ♂ | $RI_n$ | 27.27 | 57.50 | 0 | 97.85 | 100 | 100 |
| | $C_n$ | 27.27 | 41.82 | 0 | 30.25 | 0.66 | |
| Ri ♀ x Gu ♂ | $RI_n$ | NT | 0 | 0 | 100 | 0 | 100 |
| | $C_n$ | NA | NA | NA | NA | NA | |
| Gi ♀ x Ru ♂ | $RI_n$ | NT | 71.18 | 0.00 | 100 | 0 | 100 |
| | $C_n$ | NA | NA | NA | NA | NA | |
| Ru ♀ x Gi ♂ | $RI_n$ | 0* | 0 | 0 | 100 | 0 | 100 |
| | $C_n$ | 0 | 0 | 0 | 100 | 0 | |
| Gu ♀ x Ri ♂ | $RI_n$ | 60.98 | 57.75 | 33.20 | 100 | 0 | 100 |
| | $C_n$ | 60.98 | 22.53 | 5.47 | 11.02 | 0 | |
| Ri ♀ x Gi ♂ | $RI_n$ | 0* | 0 | 0 | 98.94 | 100 | 100 |
| | $C_n$ | 0 | 0 | 0 | 98.94 | 1.06 | |
| Gi ♀ x Ri ♂ | $RI_n$ | 48.72 | 62.28 | 33.20 | 98.85 | 100 | 100 |
| | $C_n$ | 48.72 | 31.94 | 6.42 | 12.77 | 0.15 | |

**Disassortative mating was found in these crosses.*
